# Computational divergence analysis reveals the existence of regulatory degeneration and supports HDAC1 as a potential drug target for Alzheimer’s disease

**DOI:** 10.1101/2023.10.05.561015

**Authors:** Qun Wang, Zhenzhen Zhao, Dong Lu, Hong Xu, Jianhua Xia, Weidong Zhang, Guofeng Meng

## Abstract

Epigenetic dysregulation has been widely reported in patients of Alzheimer’s disease (AD) and epigenetic drugs are gaining particular interest as a potential candidate therapy target. However, it is less clear how epigenetic dysregulation contributes to AD development. In this work, we performed regulatory divergence analysis using large-scale AD brain RNA-seq data and reported a widespread existence of regulatory degeneration among AD patients. It seems that transcription factor (TF)-mediated regulations get weakened or lost during AD development, resulting in disruption of normal neuronal function, especially including protein degradation, neuroinflammation, mitochondria and synaptic dysfunction. The regulatory degeneration burden (RDB) is well correlated with the detrimental clinical manifestations of AD patients. Studies of epigenetic marks, including histone modification, open chromatin accessibility and three TF binding sites supported the existence of regulatory degeneration. It suggested that epigenetic dysregulation contributed to regulatory degeneration, which also explained the consequence of epigenetic dysregulation. Among the epigenetic regulators, HDAC1 was proposed as a potential participator in such a process. Overall, our computational analysis suggested a novel causal mechanism of AD development and proposed HDAC1 as a drug target to treat AD.

## 1 Introduction

Alzheimer’s disease (AD) is a heterogeneous chronic neurodegenerative disease that has been intensively studied for decades. However, its causal mechanisms remain elusive and there is also no effective drug to cure it [1, 2, 3]. Integrated systematic approaches, especially coexpression regulatory network analysis, have advanced our understanding of AD in different ways [4, 5, 6, 7, 8]. In such studies, biological networks are constructed using expression data to identify the gene modules related to AD genesis and development by integrating quantitative evaluation. In our previous work, we also observed transcriptional dysregulation in AD patients and identified a core network [9]. Combining computational prediction with experimental perturbation allows the discovery of causal pathways and regulators, which can be used for therapeutic targets. Compared to studies performed at the single gene level, network-based analysis provides more comprehensive insights into AD causal mechanisms. However, existing studies usually take less consideration of patient heterogeneity. With the advancement of the scientific community for AD studies (e.g. AMP-AD project), large cohorts have been collected for integrated analysis. Such advancements persuade us to rethink the complex dysregulation mechanisms among AD patients.

Epigenetic regulation controls transcription regulation [10, 11]. More studies have pointed out the importance of epigenetic dysregulation in AD genesis and development, especially including DNA methylation and histone acetylation [12, 13, 14, 15, 16]. For example, genome-wide investigation of DNA methylation patterns indicated decrements of DNA methylation of enhancer regions of AD patients [17, 18, 15]. Histone modification studies to H4K16ac, H3K9ac and H3K27ac suggested a decreased epigenetic regulation, which caused dysregulation of important AD risk genes [19, 20, 21]. Additionally, recent studies reported that large-scale changes in H3K27ac could be driven by tau pathology in human brains [14]. H3K4me3 and H3K27me3 get involved in epigenetic chromatin remodeling in AD [22]. In other studies, perturbation to epigenetic reader, writer and eraser indicated the potential effects of compounds targeting epigenetic regulators to rescue AD pathology [23, 24], which supports the benefits of epigenetic perturbation. Currently, epigenetic drugs are gaining particular interest as a potential candidate therapy for AD [25, 26, 27]. However, there are still gaps to understand the detailed mechanisms of epigenetic therapy. Many efforts were put into the known AD genes, such as APP, MAPT, GSK3B and other AD risk genes [28]. These findings are not enough to indicate the pharmacological mechanism, especially when AD is characterized by patient diversity and complex causal mechanisms.

In this work, we performed a systems biology study for AD patient diversity and reported that regulatory degeneration was a potential outcome of epigenetic dysregulation, which resulted in disruption of normal neuronal function, especially including protein degradation, neuroinflammation, mitochondria and synaptic dysfunction. Our findings provided new evidence for epigenetic regulators to be potential drug targets to treat AD. In detail, we utilized a bi-clustering based method to investigate regulatory divergence among a large cohort of subjects at different AD clinical stages. We found that transcription factor(TF)-mediated regulation tended to get lost in AD patients. Both clinical association and functional studies supported regulation loss closely associated with AD development and detrimental clinical outcomes. Epigenetic studies of active regulation marks, including histone modification, open chromatin accessibility and three TF binding sites, supported the existence of regulatory degeneration. Our results suggested that epigenetic dysregulation was a driver mechanism of regulatory degeneration, and HDAC1 would be a new drug target to treat AD.

## 2 Results

### 2.1 Regulatory divergence analysis indicates the widespread existence of regulation loss

We attempted to explore the divergence of transcription regulation among AD patients using large-scale transcriptomic data. Therefore, we developed a bi-clustering based algorithm, which could subset AD patients into two groups according to the existence or strength of TF-mediated regulation (see Figure 1(a) and Supplementary Methods for detailed algorithm information). The design of this algorithm is based on two assumptions: (1) that we can identify a set of biomarker genes to indicate the TF regulatory activity; and (2) that AD patients can be clustered into groups with different TF regulation statuses. As shown in Figure 1(a,b), this algorithm takes only expression data as the input and then outputs a subset of patients that share similar TF-gene regulation patterns. Active TF regulation is identified if it satisfies the following three criteria: (1) TF-gene co-expression correlation |*r*| is greater than 0.8, which is a strict cutoff to identify biomarker genes indicating TF regulatory activity; (2) the selected patient subset has more than 50 patients; (3) TF strictly regulates least 30 genes. In case no TF satisfies the cutoff of |*r*| > 0.8 in any subset of patients, the TFs will be assigned with a type of “no regulation” (NR). NR regulators are not considered in next-step studies for the unclear regulatory role. The regulatory types of other TFs are determined by their regulation strength in the remaining patients. For example, “DR” is assigned if the |*r*| of the remaining patient subset satisfies *r* > 0.6, which indicates that such a TF has a dominant regulatory role in all patients. “WR” and “MR” indicate a weakened or missed regulation when |*r*| is above or below 0.3. Patients with weakened or missed regulation are supposed to have regulation loss or regulatory degeneration.

**Figure 1:**
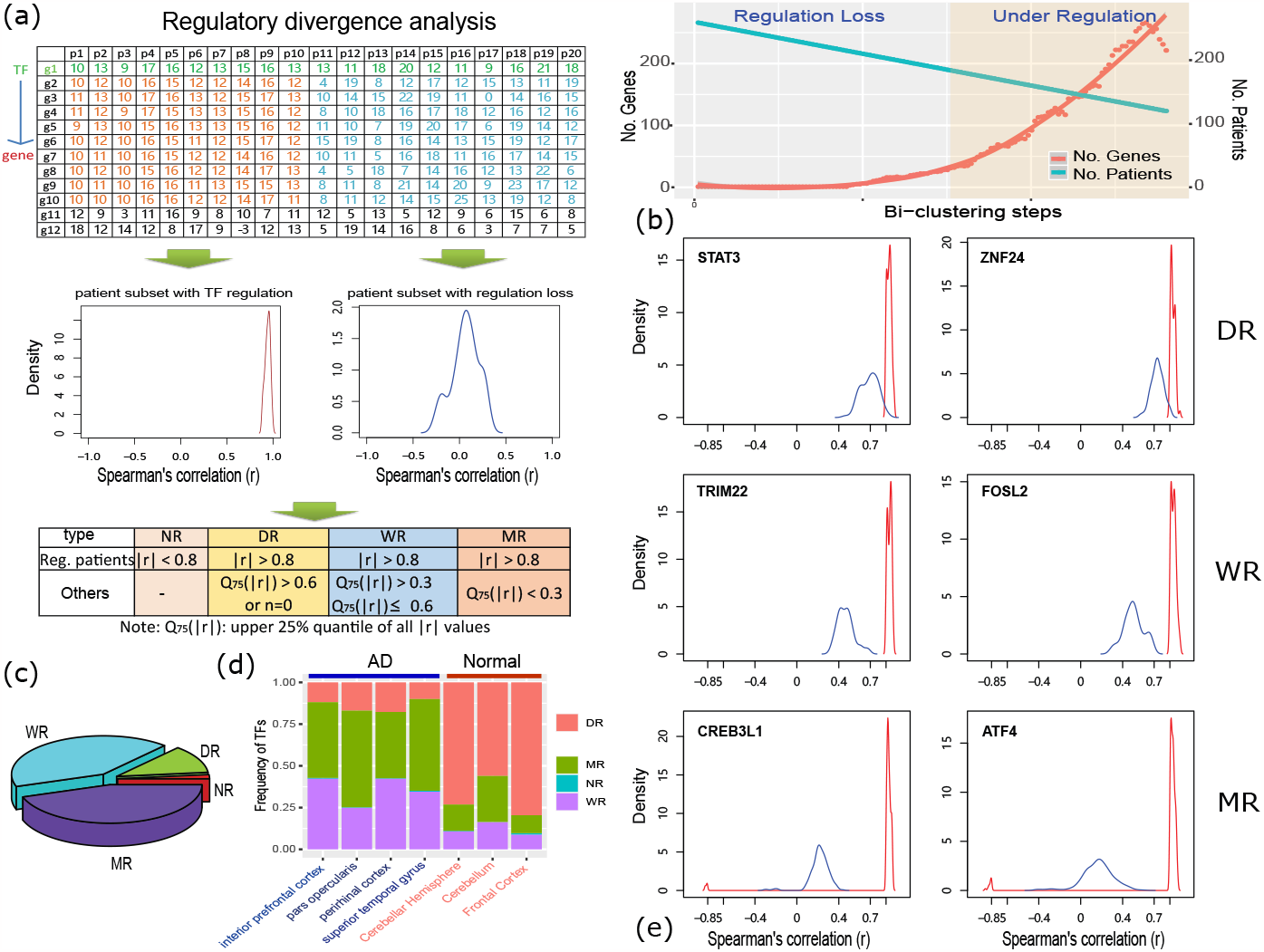
Regulatory divergence analysis indicates the widespread existence of regulation loss. (a) An analysis method to explore regulatory divergence using large-scale RNA-seq data of AD patients. In this process, a bi-clustering algorithm is applied to cluster patient samples into two groups, which may have different TF-mediated regulation statuses. Based on the strength of regulatory loss, TF-mediated regulation is assigned as one of four types: no regulation (NR), dominant regulation (DR), weakened regulation (WR) or missed regulation(MR); (b) the dynamic curve of gene and subject number during the bi-clustering analysis, reflecting the regulatory activity of studied TFs; (c) the distribution of predicted regulation types, where MR and WR are most observed in AD samples; (d) MR and WR are often observed in brain regions of AD patients but less in normal brain tissues. Here, expression data of normal subjects in three brain regions, including cerebellar hemisphere, cerebellum and frontal cortex, were collected from GTEx database and this is used to evaluate the robustness of our algorithm; (e) TF-gene correlation distribution of 6 exemplary regulators, where TFs take dominantly regulatory roles in some subjects (red line) while their regulations are weakened or missed in other patients (blue line), indicating the existence of regulation loss among AD patients.

Before application, we performed four evaluations on its reliability, including (1) the ability to identify a subset of patients with the same regulation statuses; (2) false positive ratio of bi-clustering prediction; (3) the impacts of different correlation cutoffs on the analysis results; and (4) evaluation using independent normal brain tissues. Our evaluations suggested that the predicted regulation loss was not due to technical biases of strict cutoffs and that bi-clustering analysis could recover the true regulation loss of AD patients (see details in Supplementary Results). Meanwhile, we also found that |*r*| > 0.8 was a reasonable cutoff to achieve a good analysis power for the data used in this work.

We analyzed the RNA-seq expression data for 945 autopsied samples of 364 individuals in four brain regions: the anterior prefrontal cortex (BM10), the superior temporal gyrus (BM22), the perirhinal cortex (BM36), and the pars opercularis (BM44). These subjects had diverse clinical manifestations, e.g. cognitive score and braak stages (see Figure S1 for detailed patient information). Among them, approximately 61% were diagnosed as having pathological AD or probable AD [29]. Specifically, 869 brain-expressed TFs were selected to study their regulatory status among these subjects. Among them, only nine TFs, including STAT3, ST18, CSRNP3, LMO4, CNOT7, SLC30A9, PEG3, SUB1 and MEF2C, were identified to take strict regulation in all the subjects of four brain regions at a cutoff of |*r*| > 0.8. To reduce the false negative discovery due to an excessively strict cutoff, we gradually loosed the correlation cutoff to the 75% quartile of |*r*|, *Q*_75%_(|*r*|) = 0.6 or the minimum number of co-regulated genes to 5. More regulators were identified and they were assigned with regulatory types of “DR”. For other TFs, the bi-clustering algorithm was optimized for the maximum number of genes that satisfied a cutoff of *r* > 0.8. According to coexpression correlation, we found that approximately 40% of TF had weakened regulatory roles in the remaining subjects (0.3 < *Q*_75%_(|*r*|) < 0.6) (see Figure 1(c)). Therefore, these TFs were assigned with the regulatory types of “WR”. Meanwhile, another 40% of TFs were assigned with the regulatory type of “MR”, where *Q*_75%_(|*r*|) was less than 0.3 in the remaining subjects. This result suggested that regulation loss widely existed for studied TFs. Next, we checked if regulatory loss only happened in some subset of AD patients. By investigating frequency, we found that the regulation loss was not specific to any subset of AD patients but widely existed in all subjects, and that any subjects could be under missed or weakened regulation of multiple TFs (See Figure S2). To evaluate if regulation loss is due to algorithm’s bias, we did three similar analyses using expression data of health brain samples collected from GTEx database. Dislike the analysis results of AD data, we observed more dominant regulation and less regulation loss in all three normal brain regions (see Figure 1(d)). These results indicated that regulation loss is not due to algorithm bias, and also suggested a potential relationship between regulation loss and AD genesis. Figure 1(e) showed the regulatory status of some exemplary TFs, where diverse regulatory status was observed among studied subjects.

To check if the predicted TF-regulated genes by our algorithm are really regulated by TFs, we performed TF binding motif over-representation analysis in the promoter sequences of these genes. Using the annotation of RcisTarget [30], we selected 487 TFs for evaluation and found that 31% of them were enriched with corresponding TF binding motifs (see Table S1), which was comparable to our previous findings [32]. This result indicated that TF-gene regulation identified by bi-clustering analysis was more bound by predicted TFs. We speculated that the decreased gene expression of TF genes contributed to the regulation loss. Hence, we evaluated the differential expression statuses of WR and MR TFs and found that only about 10%-15% of TF genes displayed significant expression differences between subjects with or without regulation loss at a cutoff of *p* < 0.01 (see Table S2), suggesting that most of the regulation loss was not related to decreased expression of TF genes. We further explored the transcript isoform usage of undifferentially expressed regulators using RNA-seq data [33]. However, we failed to find a strong switch in isoform usage, especially for the abundantly expressed isoforms. Overall, it seems that regulation loss is beyond expression changes of regulator genes.

In summary, we developed a bi-clustering-based method to study the regulatory divergence of AD patients and reported a widespread existence of TF-mediated regulation loss; extensive evaluation suggested that regulation loss was not due to algorithm biases or differential expression of TF genes.

### 2.2 Regulation loss contributes to disease development of AD

We investigated the disease relevance of regulation loss with AD-related clinical traits. In this step, patients were clustered by our algorithm into two non-overlapping groups: the ones under TF regulation and the ones with regulation loss. Then, two groups were checked for the value difference of clinical traits, including cognitive score (CDR), Braak score (braak) and amyloid plaque mean size (plaque), by Kolmogorov–Smirnov (K-S) test. The TFs with a significant difference were supposed to be associated with AD-related clinical traits. To ensure confidence, we randomly shuffled clinical trait values of AD patients and repeated the same analysis to estimate the false discovery ratio (FDR). At a cutoff of *p* < 0.01 and FDR < 0.05, 291, 277, 335 and 159 TFs were predicted to be associated with at least one of three clinical traits, accounting for about 38%, 36%, 47%, and 22% of MR/WR TFs in four brain regions, respectively (see Figure S3 and Table S3). To further evaluate the validity of these observations, we performed another round of simulation evaluation by repeating the same analysis using random sample combinations. Under this setting, the maximum number of significant TFs was less than 10 for all three clinical traits and the possibility for our observation was nearly impossible (*p* = 0), which suggested a good confidence of the association between regulation loss and clinical traits.

Figure 2(a) shows some exemplary TFs and their clinical associations. As described above, two subsets of patients showed significant value differences of clinical traits. By searching the published literature, we found that many of these TFs had been reported for AD- or brain-related function. For example, OLIG2 (Oligodendrocyte transcription factor) contributes to cognitive defects of Down syndrome [34]. Evidence suggests that OLIG2 participates in fate switch of neurons in AD patients [35]. In our result, OLIG2 was an MR regulator and significantly associated with both CDR (*p* = 6.86*e* − 7) and Plague (*p* = 7.19*e* − 9) in three brain regions, including BM10, BM22 and BM44. THRA (Thyroid Hormone Receptor, Alpha) is an MR regulator associated with CDR (*p* = 2.02*e* − 8) and Plague (*p* = 1.04*e* − 6) in two brain regions. It is essential for normal neural development and regeneration [36]. THRA has been reported to have a weak genetic link with AD [37]. ATF4 has been widely reported for the increased expression in AD patients and transcriptional mediator roles in neuron degeneration, metabolism and memory formation [38].

**Figure 2:**
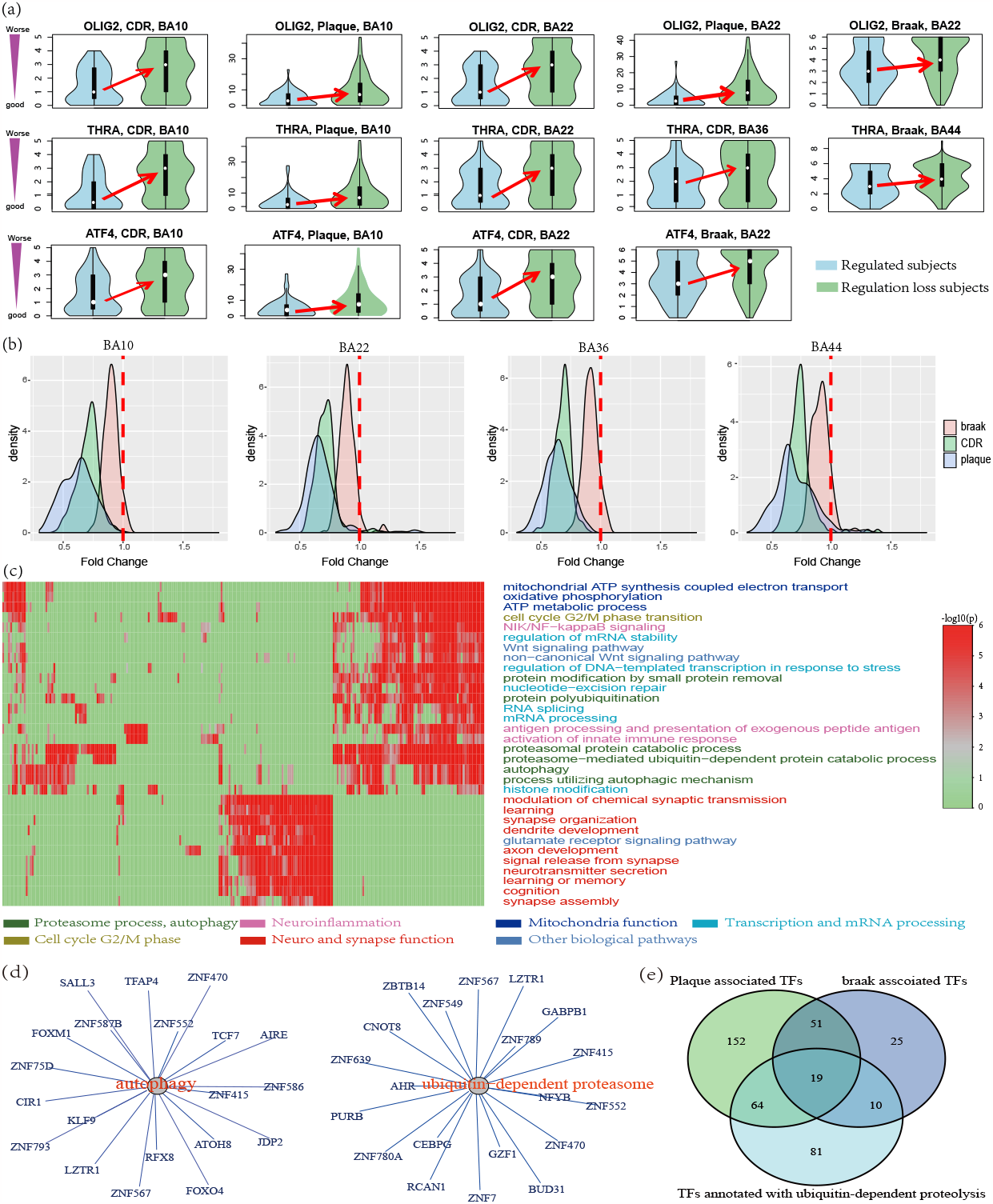
TF-mediated regulation loss contributes to AD development. (a) Clinical association of the regulation loss mediated by some exemplary TFs. Here, the patients with or without regulation loss showed a significant differences in clinical manifestation. (b) Regulation loss was always associated with detrimental clinical outcomes. In this study, about 250 TFs were identified to be associated with at least one AD-related clinical trait in each brain region; fold change indicated the clinical trait value division in the patients with or without regulation loss. Lower fold changes (< 1) indicate changes to the more detrimental clinical outcomes. (c) Functional involvement of regulation lost TFs was annotated by functional enrichment analysis of TF-regulated genes. The enriched terms included protein degradation, neuroinflammation, mitochondrial function, neuronal and synaptic functions. (d) The TFs annotated with functional involvement of ubiquitin-dependent proteasome and autophagy process. (e) The TFs annotated with ubiquitin-dependent proteolysis were more associated with plague and braak scores (*p* = 0.005).

From these examples, we also observed consistent association direction, where TF-mediated regulation losses was more observed in the patients with a detrimental clinical manifestation (see Figure 2(a)). We checked the association direction of other TFs by calculating the fold change of clinical trait values. As shown in Figure 2(b), nearly all TF-mediated regulation loss were associated with detrimental clinical outcomes, such as declined cognitive ability, increased Braak stage and increased amyloid plaque size. A similar tendency was consistently observed in four brain regions. Among the three clinical traits, CDR and plague always had a stronger clinical association, e.g. more associated TFs or larger fold changes. It seems that TF-mediated regulation loss is associated with severe AD stages.

In the bi-clustering analysis, our algorithm identified the TF-regulated genes. In total, there were 6300 such genes that regulated by at least one of MR and WR TFs. Among them, 43% of genes were regulated by only one TF and 80% were regulated by less than 5 TFs (see Figure S4). To understand the consequence of regulation loss, we performed functional enrichment analysis using TF-regulated genes. Figure 2(c) highlights the pathways and biological processes associated with regulation loss. Protein degradation, especially the ubiquitin-proteasome system (UPS), was the most affected biological process (see Figure S5), accounting for more than 50% of TFs in all four brain regions, consistent with the growing evidence for a tight link between UPS impairment and abnormal aggregation of toxic proteins [39, 40]. Additionally, autophagy, another biological process of protein degradation, was enriched in the regulated genes of more than 60 TFs. Figure 2(d) shows the UPS and autophagy associated TFs. Next, we checked if the expression of these TFs was correlated with plaque and braak scores. We found that more than half of the TFs annotated with ubiquitin-dependent proteasome function were also associated with plaque and braak scores in gene expression level (*p* < 0.005) (see Figure 2(e) and Figure S6). These results indicated a close association between regulation loss and protein degradation process.

Neuroinflammation was the second most enriched biological process. Terms, such as “NIK/NF-kappaB signaling” and “innate immune response activating cell surface receptor signaling pathway” are enriched with about 40% of TFs in all four brain regions. This finding is consistent with the reports that activation of inflammation contributes to AD pathogenesis [41, 42]. Mitochondria-related functions, such as “electron transport chain” and “oxidative phosphorylation”, were enriched in about 40% of TFs. This is supported by reports for the role of mitochondrial dysfunction and oxidative damage in the genesis of AD [43, 44]. Neuronal and synaptic function related processes were enriched with about 20% of TFs [45]. Other enriched processes include Wnt signalling pathway, rRNA processing, histone modification and other processes.

Overall, our results suggest that regulation loss is associated with detrimental clinical manifestation of AD patients and also contributes to AD development by affecting AD-related pathways, especially including protein degradation and Neuroinflammation.

### 2.3 Regulatory degeneration burden better correlates with AD clinical outcomes

We have identified the regulation loss for about 200 TFs, all of which possibly contribute to AD development. To understand the accumulated effects of these TFs, we introduced a systematic measurement, regulatory degeneration burden (RDB), which accounts for an accumulated degree of regulation loss (see detailed description in Methods section). In this way, each AD patient will be assigned an RDB score to indicate his/her degree of regulation degeneration.

We found that RDB had good correlations with all three clinical traits, especially including CDR and Plague (see Figure 3(a)). In all four brain regions, RDB had the best correlation with CDR, where Spearman’s correlation was up to *r* = 0.52 in BM10. In BM10, RDB had the best correlation with plaque score at *r* = 0.50. In other brain regions, RDB also achieved good correlations with CDR and Plaque, too. As an evaluation, we also investigated the relevance with other covariates, such as age and sex. As shown in Figure 3(b), RDB only exhibited a strong correlation with AD-related clinical traits. To check if neuron loss led to RDB, we performed cellular deconvolution using bulk RNA-seq data and calculated the cellular composition of six brain cell types, including oli (oligodendrocytes), mic (microglia), ast (astrocytes), opc (oligodendrocyte progenitors), in (inhibitory neurons), ex (excitatory neurons). As showed in Figure S7, the composition of multiple cell types was weakly correlated with RDB. Considering that cellular composition was also associated with clinical manifestation (see Figure S7), we performed partial correlations to remove the contribution of clinical traits. The new correlation results did not support any association between RDB and cellular composition (see Figure 3(c)). This result suggested that RDB was not caused by neuron composition difference or neuron loss.

**Figure 3:**
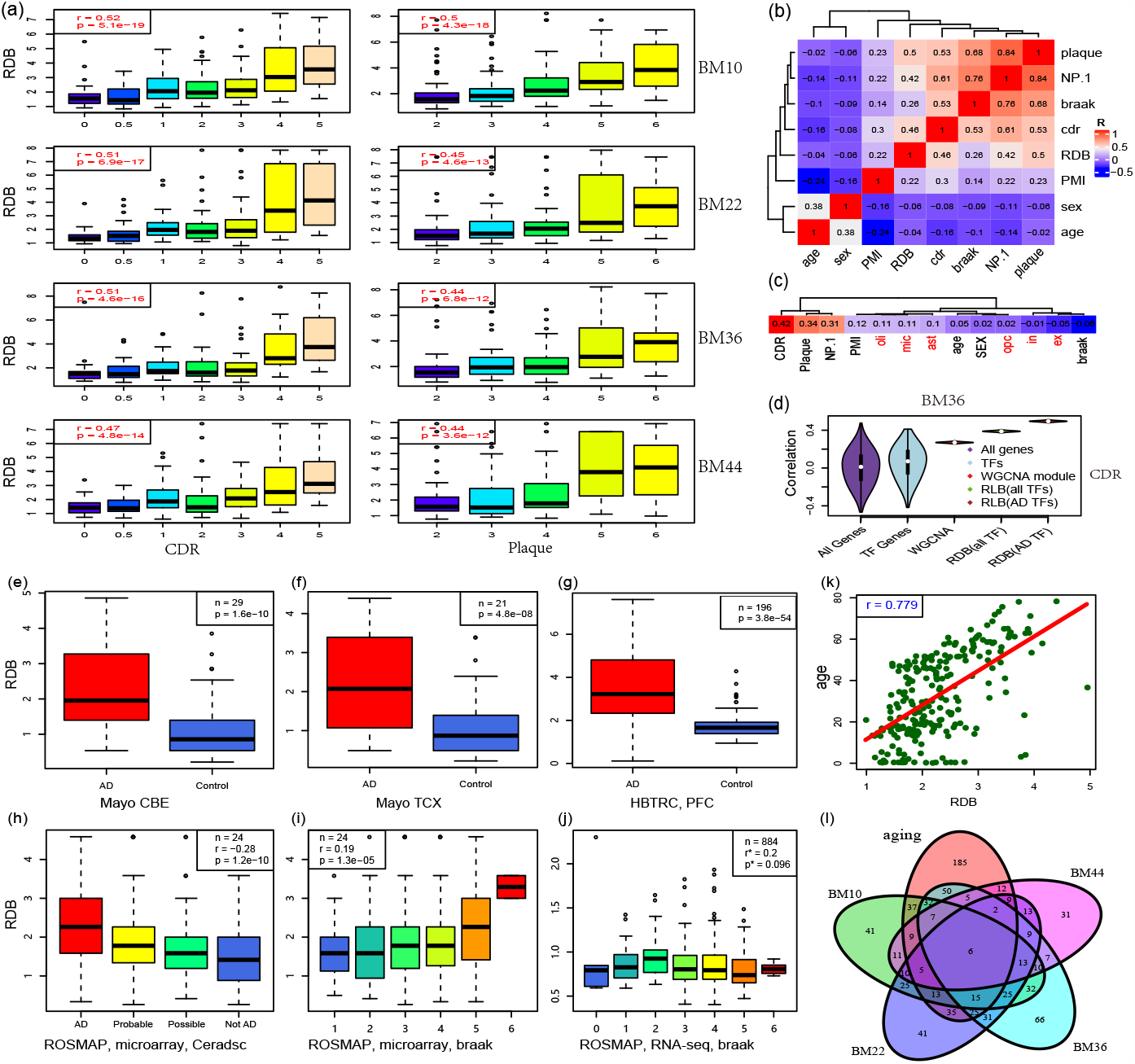
Regulatory degeneration burden better correlates with AD clinical outcomes. (a) Association between regulatory degeneration burden and clinical outcomes in four brain regions, where the correlation value is up to *r* = 0.5; (b) Regulatory degeneration burden is only associated with AD-related clinical traits. PMI: Post-Mortem Interval; NP.1: Neuropathology Category. (c) The cell type compositions of brain samples were not associated with regulatory degeneration burden. Here, the brain samples were deconvoluted into six cell types: oli (oligodendrocytes), mic (microglia), ast (astrocytes), opc (oligo-dendrocyte progenitors), in (inhibitory neurons), ex (excitatory neurons), and partial correlation analysis did not indicate any association between RDB and neuron loss. (d) Regulatory degeneration burden better indicates AD development than genes, transcription factors and WGCNA modules. (e-j) showed Regulatory degeneration analysis in five independent datasets, including (e) Mayo CBE, (f) Mayo TCX, (g) HBTRC PFC, (h-i) ROSMAP microarray and (j) ROSMAP RNA-seq. (k) Regulatory degeneration burden well correlates with ages, indicating that regulatory degeneration contributes to the ageing process. (l) TF overlaps among the ageing process and AD in four brain regions, which suggests the close relationship between AD and the ageing process.

Transcriptomic features, e.g. gene expression of some biomarker genes, have been previously explored for their associations with clinical features [6]. However, most of these studies failed to identify genes or modules with strong correlations to AD clinical traits. We compared RDB with these measurements (see Figure 3(d) and Figure S8). At the single gene level, RDB showed better performance in BM10, BM22 and BM44 regions. In BM36, there were only 34 genes that had a better correlation with CDR or Plaque than RDB. Among them, the maximum correlation was observed with CDR for C4B at *r* = 0.55. Among the TF genes, there are only 2 TFs showed a better correlation with CDR or Plaque than RDB, including HDAC1(r=0.54) and ZNF215 (r=0.52). Next, we performed WGCNA network analysis in each brain region and the predicted modules were evaluated for all clinical traits. Without including the grey module, the maximum correlation value (*r* = 0.34) was observed between CDR and a module predicted in the BM36 region, which was lower than that of RDB. Our evaluation suggests that RDB better correlates with AD clinical outcomes than the measurements using single gene or gene modules.

To evaluate the confidence of our findings, we repeated the same analysis using five independent expression data collected from the published studies. They included ROSMAP expression data using microarray [4] and RNA-seq [5], HBTRC microarray study [4], Mayo’s RNAseq study for cerebellum (CBE) and temporal cortex (TCX) [46]. The dataset from Mayo and HBTRC had no extra clinical annotations and only a binary disease status was available. Thus, RDB values were checked between the AD patients and control subjects. As shown in Figure 3(e-g), AD patients displayed more regulatory degeneration than control subjects. Among them, HBTRC dataset had more subjects (463 subjects) and identified 196 AD-associated TFs. Statistical test suggested a significant regulatory degeneration in AD patients (*p* = 3.8*e* − 54). ROSMAP dataset included both microarray and RNA-seq data. Analysis results using the microarray data indicated a positive correlation between an increased RDB value and detrimental clinical outcome (see Figure 3(h, i)). However, using ROSMAP RNA-seq data, RDB values did not indicate a consistent tendency. As shown in Figure 3(j), a positive correlation was observed when Braak score was less 3 and a negatively correlated tendency was observed when Braak was greater than 3. It seems that the ROSMAP RNA-seq dataset only supports the hypothesized relationship between RDB and clinical outcomes at an early stage of AD. We evaluated ROSMAP RNA-seq data but did not find any clue for such inconsistency. Considering the analysis results using microarray, we still believe regulatory degeneration to be associated with AD clinical outcomes. Overall, five independent datasets partially or completely support the same conclusion that regulatory degeneration correlates with the detrimental clinical outcomes of AD patients.

Age is the biggest risk for AD. Therefore, we performed regulatory degeneration analysis using microarray expression data from human brain prefrontal cortex. This dataset included 229 AD-free participants with a broad age range from 0 to 70 years old [47]. Using the same analysis parameter, 447 regulators were predicted to be associated with ages. Like that of AD, the ageing process was accompanied by increased RDB, where Spearman’s correlation between RDB and ages is 0.779 (see Figure 3(k)), which indicated the important roles of regulation loss in the ageing process. We analyzed the overlap of regulation-lost regulators from AD and ageing and found that 262 TFs were shared (see Figure 3(l)). However, this overlap was not statistically significant (*p* = 0.18). This result suggests that the regulation degeneration in AD patients is not the exact same regulation degeneration that occurs during the ageing process. This was also supported by our observation that RDB values were not correlated with ages of AD patients (Figure 3(b)).

In summary, our results indicated that regulatory degeneration burden better predicted AD clinical outcomes, which was repeatedly observed in multiple independent datasets. Meanwhile, regulatory degeneration burden also contributed to the ageing process, where the correlation with ages was up to 0.779.

### 2.4 Active Epigenetic regulation loss is a potential driver of regulatory degeneration

To confirm the existence of regulation loss, we performed ChIP-seq analysis to identify the binding sites of TFs. Three TFs were selected, including ATF4, OLIG2 and THRA. We used two strategies to study their binding statuses. The first option is to do differential peaking analysis to study the peak intensity between AD and normal samples. Another one is to count the absolute number of peaks among the AD patients and normal samples. In Figure 4(a), we showed the result of differential peaking analysis to Olig2 binding sites and found that the ratio of binding site gain against loss was 1.19 : 1, which did not support regulation loss. Then, we counted the number of Olig2 binding sites for each sample and observed clear Olig2 binding site loss in AD patients. As shown in Figure 4(b),7539 peaks were missed in AD patients while only 874 peaks were gained. This result suggested that Olig2 binding sites were significantly lost in AD patients. We performed the same analysis to ATF4 and THRA. Unlike Olig2, we only observed a weak ATF4 binding peak missing in AD patients. However, a stronger tendency of binding intensity weakening was observed in differential peaking analysis for both TFs, especially for THRA (see Figure 4(i-l)). We also analyzed the overlaps between lost TF binding sites and the corresponding dysregulated genes. We mapped ChIP-seq TF peaks to their nearby genes (see Figure S9). We observed significant overlaps with the lost binding sites of ATF4 (*p* = 0.01) and THRA (*p* = 0.001) (see Figure S10), which supports the contribution of lost TF binding sites. It has been reported for the benefits of epigenetic perturbation to rescue AD-related clinical manifestations [48, 49]. Among them, histone deacetylase inhibition has been reported for clear benefits after their inhibition [23]. Therefore, we investigated if epigenetic dysregulation can be the driver of transcription regulatory degeneration. H3K27ac has been observed to display differential enrichment in AD patients, especially in the regulatory regions of AD risk genes [20]. We performed peak calling analysis on H3K27ac ChIP-seq data and found fewer peaks in AD patients compared to that of controls, suggesting an active H3K27ac regulation loss in AD patients (see Figure 4(a)). Applying a modified analysis pipeline originally introduced in [20], we re-evaluated H3K27ac enriched regions and observed a strong tendency of H3K27ac mark loss in AD patients, where the ratio of loss against gain was 1 : 0.37 (see Figure 4(b)). Then, we checked if the H3K27ac loss was associated with dysregulated gene expression. Using the downstream target genes of the top 10 dysregulated TFs (see Table S3), a significant overlap was observed between dysregulated genes and the genes with H3K27ac loss in the promoter regions (*p* = 5.08*e* – 32 by Fisher’s exact test) (see Figure 4(c). Similar but weak results were observed with H4K16ac, which is another active regulation mark [19] (see Figure S11). Next, we performed the assay for transposase-accessible chromatin using sequencing (ATAC-seq) to identify active regulatory regions. At a cutoff of *q*-value < 0.05, we found 207,765 open chromatin regions in 12 subjects. Both AD patients and control subjects displayed diversity in ATAC-seq peaks and there was only a weak tendency of open chromatin region loss in AD patients (*p* = 0.3) (see Figure 4(d)). We performed differential peaking analysis and found a significant tendency of open chromatin region weakening in AD patients, where the ratio of loss again gain is 1 : 0.46 (see Figure 4(e)), which supported open chromatin region loss in AD patients. We then evaluated their association with dysregulated gene expression. Using the genes mentioned above, we observed a significant overlap (*p* = 8.05*e* − 9)(see Figure 4(f)), indicating that lost open chromatin accessible regions were significantly related to dysregulated gene expression.

**Figure 4:**
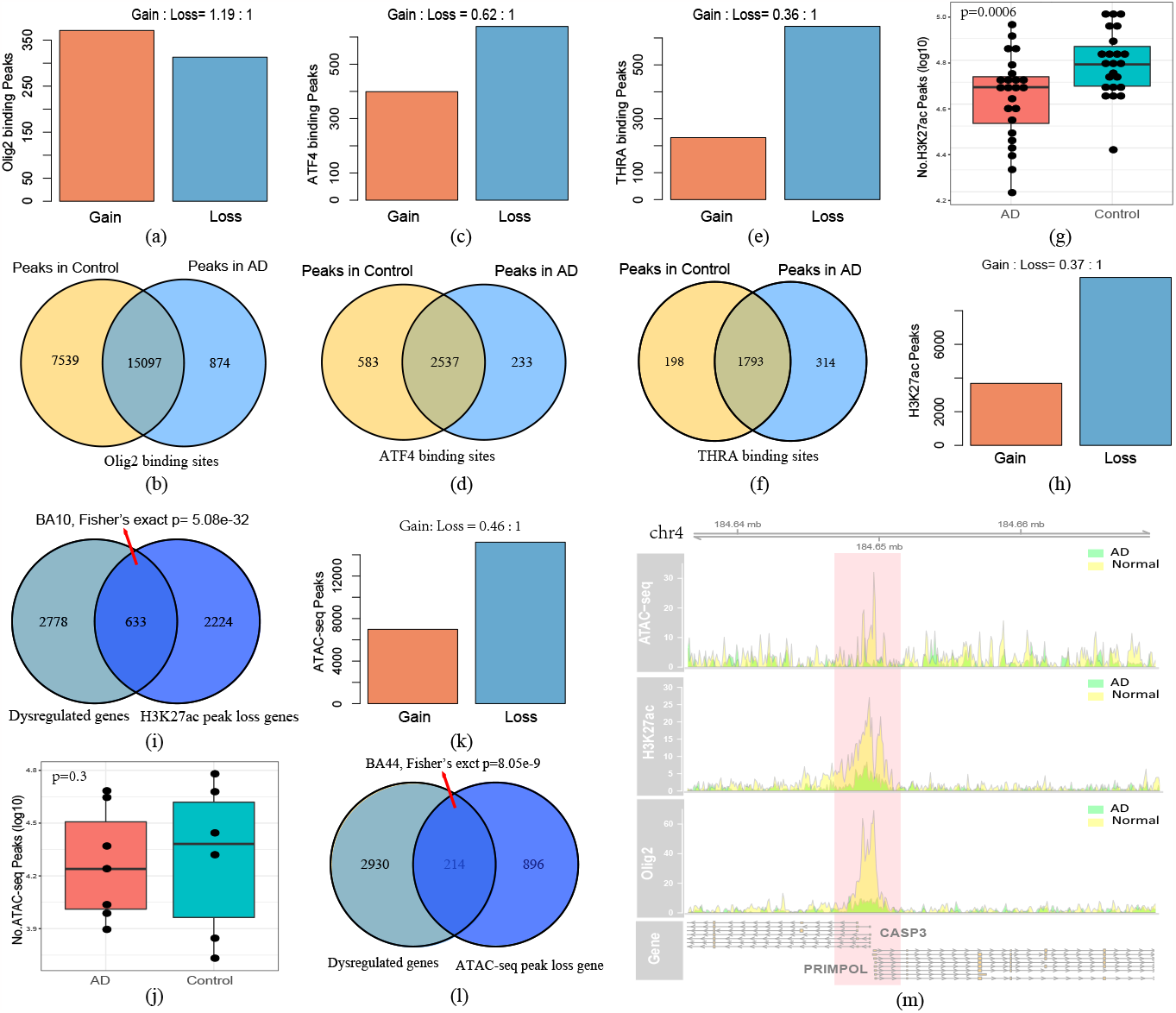
Active Epigenetic marker loss is a potential driver of regulatory degeneration. The active regulation marks are checked by differential peaking analysis and the number of peaks. (a-b) the analysis results of Olig2 binding sites, where Olig2 binding site loss is observed in AD patients but not binding intensity; (c-f) the analysis results to ATF4 and THRA binding sites, where they have a tendency of decreased TF binding intensity in AD patient; (g) A tendency of H3K27ac mark loss in AD patients is observed based on the peak calling analysis (*p* = 0.0006 by t-test), indicating lost activity of transcription regulation; (h) differential peaking analysis also suggests a tendency of H3K27ac mark loss in AD patients; (i) the genes with lost H3K27ac histone modification are significantly overlapped with the dysregulated genes that are identified in bi-clustering analysis; (j) ATAC-seq peaks have a weak tendency of ATAC peak loss in AD patients (*p* = 0.3); (k) differential peaking analysis to ATAC-seq data suggests more chromatin accessibility loss in AD patients; (l) the genes associated with chromatin accessibility loss are significantly overlapped with the AD dysregulated genes; (m) One region with the loss of open chromatin accessible region, H3K27ac mark and Olig binding site in AD patients.

In summary, our results suggested that there was active epigenetic regulation loss in AD patients, and they could be a potential driver of regulatory degeneration.

### 2.5 HDAC1 contributes to regulatory degeneration of AD

To evaluate the AD association of histone deacetylase, we performed differential expression analysis and clinical association studies for members of HDAC family. Figure S12 showed the analysis results using MSBB data. We found that most of HDACs had good associations with AD at transcriptional level, including significant differential expression and good correlation with clinical features. Among them, HDAC1 displayed the best association with AD, where the correlation with cognitive scores was *r* = 0.53 in the perirhinal cortex regions (BM36) (see Figure 5(a)). In other brain regions, HDAC1 also displayed best AD association with other clinical manifestations, e.g. Braak score (see Figure S12). This result proposed that HDAC1 potentially contributed to regulatory degeneration in AD.

**Figure 5:**
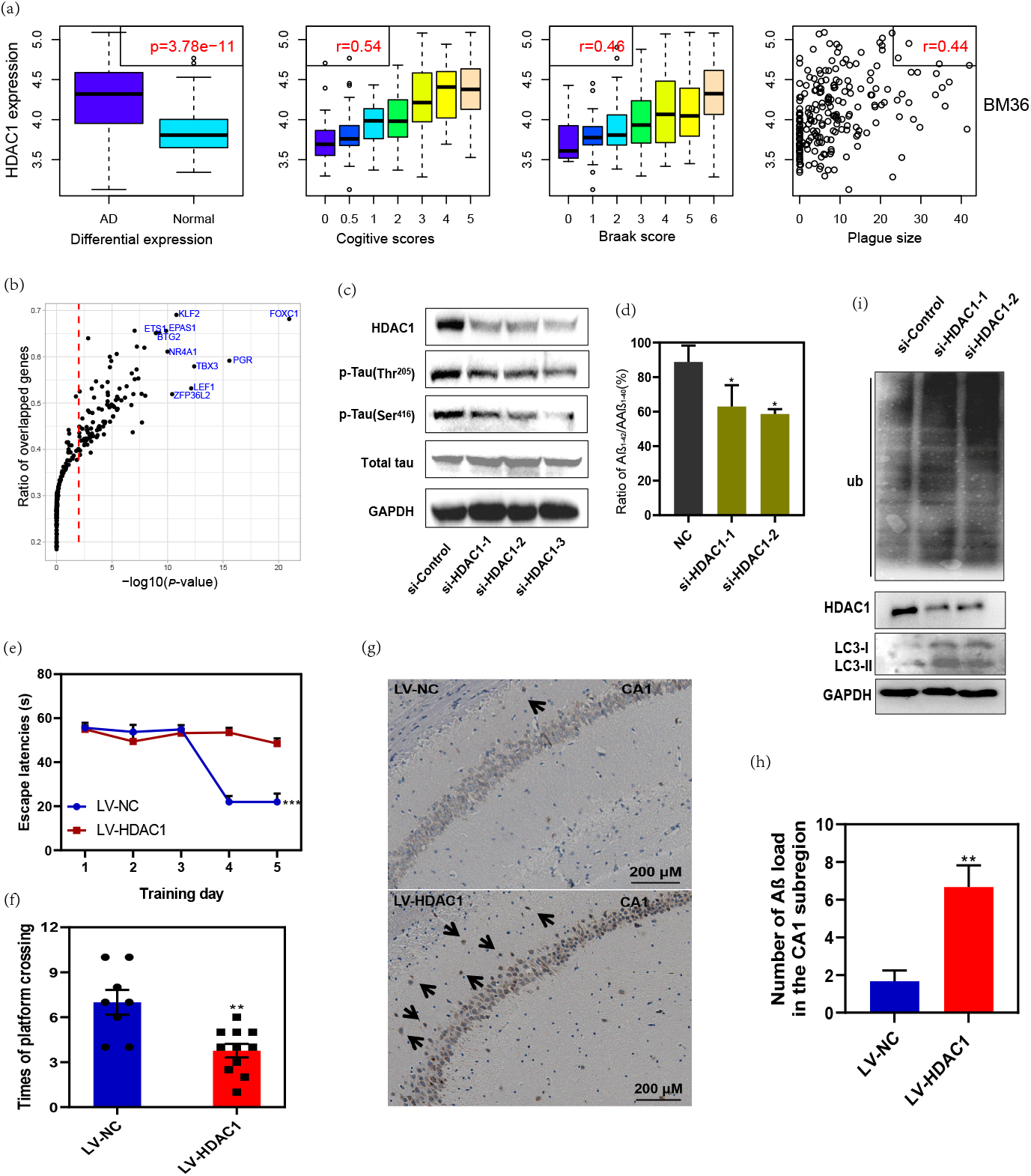
HDAC1 contributes regulatory degeneration in AD. (a) HDAC1 gene is differentially expressed in AD samples and its expression is well correlated with AD-related clinical features. (b) The differential expressed genes induced by HDAC1 over-expression were significantly overlapped with the genes affected by regulatory degeneration. (c) HDAC1 silencing is associated with lower Tau phosphorylation levels at Thr^205^ and Ser^416^ sites; (d) HDAC1 silencing associated with a lower Aβ_1−42_:Aβ_1−40_ Ratio. The results of water mirror maze test, including (e) longer time of escape latency and (f) decreased number of platform crossing were observed in the water mirror maze studies to HDAC1 over-expressed transgenic mice. (g) and (h) The deposition of Aβ in HDAC1 over-expressed transgenic mice was significantly increased than control animals.(i) HDAC1 silencing is associated with higher protein ubiquitin levels and increased autophagic activity in cells.

To validate this hypothesis, we performed RNA-seq study on the HDAC1 over-expressed SH-SY5Y cells and identified 5059 differentially expressed genes (DEGs) at a cutoff of adjusted-*p* < 0.01. Then, we checked if HDAC1 over-expression contributed to TF-mediated regulatory degeneration by checking the overlaps between DEGs induced by HDAC1 over-expression and the genes affected by TF-mediated regulatory degeneration. We found that the genes regulated by 291 TFs, accounting for 43.3% of the studied TFs, are significantly enriched with HDAC1-induced DEGs at a cutoff of *p* < 0.01 by Fisher’s exact test (see Figure 5(b)). To validate the confidence of such observation, we performed simulation studies by assigning random genes as the DEGs after HDAC1 over-expression, and found that downstream genes of only 1.3% of TFs are enriched with HDAC1 over-expression-induced DEGs. This result supported a hypothesis that HDAC1 contributed to regulatory degeneration.

The two hallmark pathologies of AD are the extracellular plaque deposits of the β-amyloid peptide (Aβ) and the flame-shaped neurofibrillary tangles of the microtubule-binding protein tau [50]. Excessive tau phosphorylation is associated with AD-related neurofibrillary tangles [51]. Therefore, we checked the phosphorylation levels of tau proteins and found that the Thr^205^ and Ser^416^ sites showed a remarkably decreased phosphorylation level after genetic silencing of HDAC1 compared to the control group (see Figure 5(c)). We also examined the ratio of neurotoxic Aβ_1−42_ and nontoxic Aβ_1−40_ after HDAC1 silencing. As expected, we observed a decrease in neurotoxic Aβ_1−42_ and an increase in nontoxic Aβ_1−40_, resulting in a significantly decreased ratio of Aβ_1−42_to Aβ_1−40_ in HDAC1-silenced cells (see Figure5(d)).

To further investigate if HDAC1 contributes to AD development, we constructed HDAC1 over-expressed mouse CAG-LSL-HDAC1-3*FLAG-PolyA and performed water mirrors maze(MWM) test at 10 month of age. A one-way ANOVA was conducted to compare the spatial memory differences with normal mice. As showed in Figure 5(e), the transgenic mouse showed reduced escape latency during the training period, especially on the fourth and fifth day (groups: F(1, 20) = 26.94, P = 0.0003; days: F(3.2, 58.7) = 27.33, P < 0.001; group*×*day: F(4, 74) = 22.25, P < 0.001). During the probe trial, the times of target platform crossings of transgenic mice were decreased significantly (P = 0.036; see Figure 5(f)). Moreover, the spent time and swimming distance in the target quadrant were weakly differential in transgenic mice (P = 0.15 and P = 0.05, respectively; Figure S13(a,b)). We also examined the deposition of Aβ in the mouse hippocampus (CA1) by immunohistochemistry and found that the deposition of Aβ in HDAC1 over-expressed transgenic mice was significantly increased (*p* = 0.0026, see Figure 5(g,h)).

In the above section, we computationally predicted that ubiquitin-dependent proteolysis was the most affected process by regulatory degeneration. Therefore, we investigated if HDAC1 was also related to such a process. The overall ubiquitin levels were examined after silencing HDAC1 gene and observed a significantly increased ubiquitin level in HEK/APPsw cells (see Figure 5(i)). This result suggested HDAC1 activity was closely associated with ubiquitin-dependent proteolysis. Autophagy is another important process affected by regulatory degeneration. Therefore, we examined the level of LC3-II and found a significant increase in LC3-II expression after HDAC1 silencing (see Figure 5(i)). This result indicated that autophagy was also elevated in the cells. Combined with the above findings, these results suggested that HDAC1 was associated with regulatory degeneration in patients mainly by affecting ubiquitin-dependent proteasome and autophagy process.

## 3 Discussion

In this work, we performed the first investigation of AD patient diversity in both regulation status and clinical manifestations. We found that accumulated degeneration of transcription regulation widely existed in AD patients and contributed to the disease development, the detrimental clinical outcomes and the ageing process. This conclusion is drawn based on results using computational modeling analysis of large-scale RNA-seq data and genome-wide studies of active epigenetic marks, where we found that (1) transcriptional regulation tends to get lost in AD patients; (2) transcription regulation loss almost indicated detrimental clinical outcomes;(3) regulatory degeneration burden was better correlated with the clinical outcomes than the existing methods; (4) regulatory degeneration mainly disrupts the AD-related biological processes, especially protein degradation, neuroinflammation, mitochondrial function and synaptic function; and (5) active epigenetic regulation also get lost in AD patients.

Combining these findings, it may lead to a novel causal mechanism for AD, where brain regulation degenerates from an organized system in normal individuals into a deficient system in AD patients, which disrupts many AD-related biological processes (see Figure 6). Compared to existing hypotheses, this mechanism better integrates existing knowledge about AD under a unified framework, including the epigenetic dysregulation [12, 13, 14], disturbed gene regulatory network [4, 5, 6, 52], broad involvement of multiple biological processes, complex clinical manifestations and AD patient diversity.

**Figure 6:**
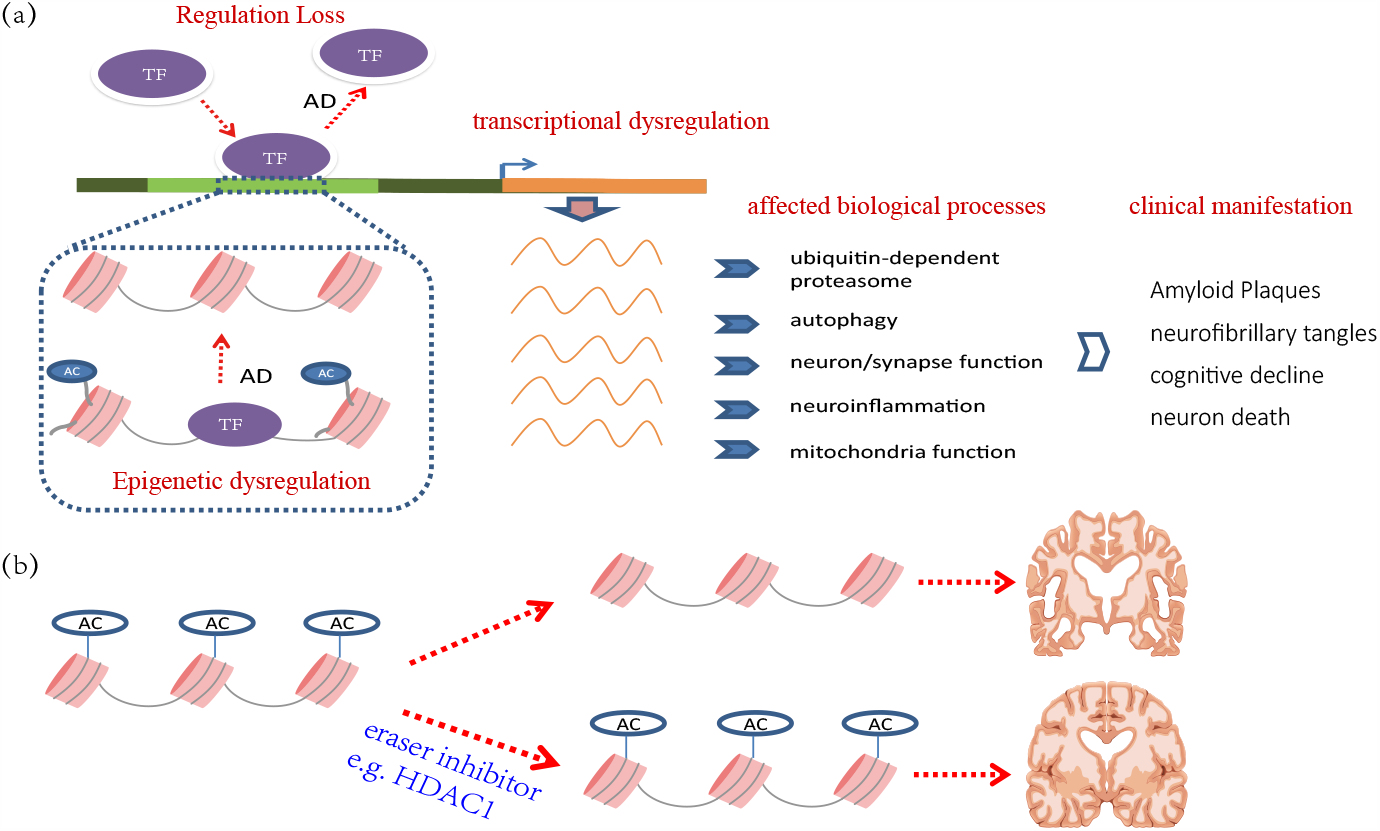
Regulatory degeneration and epigenetic dysregulation. (a) Regulatory degeneration contributes to AD genesis and development. Our results suggest that transcriptional regulation tends to get degenerated among AD patients, which disrupts the normal cellular function of brain, e.g. protein degradation, neuroinflammation, mitochondrial, neuronal/synaptic function, and contributes to detrimental clinical outcomes. This finding supports an integrative model to elaborate the causal mechanisms of AD, where brain transcriptional regulation degenerates from an organized system in normal individuals into a deficient one in AD patients. (b) Epigenetic regulators are potential therapeutic targets of AD. Both genome-wide epigenetic screening and inhibitor perturbation studies supported epigenetic regulators to be a potential drug target to cure AD; our finding reveals a potential mechanism for epigenetic drugs.

In this work, we investigated the potential mechanism of regulatory degeneration and proposed epigenetic dysregulation as a causal mechanism. Before this work, epigenetic drugs are gaining particular interest as a potential candidate therapy of AD. This is supported by genome-wide epigenetic studies to marks and perturbation studies to epigenetic writers and erasers. However, it is still not clear how such drugs affect the complex AD causal mechanisms. The importance of our finding is that it reveals a potential pharmacological mechanism of epigenetic drugs by bridging epigenetic modification to diverse AD causal mechanisms.

Among the numerous epigenetic regulators, we proposed HDAC1 as a potential participant to cause regulatory degeneration. The evidence includes that (1) H3K27ac, the catalytic substrate of HDAC1, is a differential mark in AD; (2) the expression of HDAC1 is more correlated with the clinical manifestation of AD patients. Experimental investigation also supported its roles in AD developments, by affecting the composition of β-amyloid peptide (Aβ) and phosphorylation levels of tau proteins. All these results suggested that HDAC1 was associated with AD development by affecting regulatory degeneration of AD. However, our current results cannot exclude the potential roles of other epigenetic regulators.

The current analysis pipeline has several limitations. We applied an approximating solution to identify the gene-subject combinations by gradually removing the subjects and checking the improvement of Spearman’s correlations. This algorithm might reach a local optimum or even a false solution. We used a set of arbitrary cutoffs in our bi-clustering analysis. For example, we used |*r*| > 0.8 as the cutoff of TF-gene dominant regulation and used a minimum number of 30 genes and 50 patients as the cutoff for successful bi-clustering results. Due to the limited knowledge of TF action mechanism, the selected cutoff setting may not be in accordance with the actual scenarios. Meanwhile, the bi-clustering could generate a continuous combination of subjects and genes. How to select the best gene-patient blocks is also a great challenge. We choose the combination when the number of block genes reaches the maximum values. This may not work for some TFs. In the future, we would improve the bi-clustering algorithm so that it can optimize the output based on the properties of bi-clustering output.

## 4 Data and Methods

### 4.1 Brain samples

Postmortem brain samples in the prefrontal cortex regions of 26 individuals, including 13 diagnosed with AD and 13 normal subjects were collected from the Chinese Brain Bank Center in Wuhan (CBBC, http://cbbc.scuec.edu.cn) and China Brain Bank, Zhejiang University (http://www.neuroscience.zju.edu.cn). Informed consent for autopsy has been signed for all the subjects by brain banks when the participants were in life. The clinical information of each subject was reviewed by independent neurologists with expertise in dementia and the neuropathological diagnosis was given regarding the most likely clinical diagnosis at the time of death. This study was reviewed and approved by the Ethics Committee of both brain banks and Shanghai University of Chinese Medicine for both ChIP-seq and ATAC-seq. The final sample size was limited by the available AD samples. 13 AD samples were collected and one sample was excluded for bad sample quality. We prioritized the samples with no AD or other neurological clinical manifestation as the control samples. The AD and normal samples were assigned into two groups (each size of 6): one group was used for ChIP-seq experiment of OLIG2 and ATF4 and another group was used for ChIP-seq of THRA1 and ATAC-seq. The subject information is available in Table S4.

### 4.2 Chromatin Immunoprecipitation (ChIP-seq)

The whole experiment was performed following the published protocols (see Supplementary Methods). The used the antibody includes (1) ATF4, CTS,11815S (lot#4); (2) Olig2, RnD, AF2418 (lot#UPA0718031); (3) THRA, SantaCruz, SC-56873 ((lot#J1614).

### 4.3 Assay for Transposase-Accessible Chromatin using sequencing (ATAC-seq)

ATAC-seq was performed in GENEWIZ company following the protocol introduced in [53, 54] (see Supplementary Methods).

### 4.4 Data collection and processing

The RNA-seq expression data were collected from the Mount Sinai Brain Bank (MSBB) study of Accelerating Medicines Partnership-Alzheimer’s Disease (AMP-AD) projects. The raw count data were firstly filtered to remove the non-expressed genes by setting the maximum number of subjects with counts 0 is less than 10%. Then, the count data were normalized by the TMM algorithm implemented in edgeR package [55]. A linear regression model was used to adjust the effects of covariates, including age, sex, postmortem interval (PMI) and RNA integrity number (RIN). The covariates were evaluated by the principal components to make sure their effects are excluded. The samples with obvious deviation were treated as outliers and removed for further analysis. The independent expression data, including ROSMAP study using microarray [4], ROSMAP study using RNA-seq [5], HBTRC study [4], Mayo’s RNAseq study for cerebellum (CBE) and temporal cortex (TCX) [46]. All of them can be retrieved from AMP-AD projects. Like the ROSMAP data, the RNA-seq counts data were collected and the similar processing pipeline was applied. For the microarray, the normalized data were collected and post-processing was performed like that of RNA-seq data.

### 4.5 Transcription factor

The TFs were selected based on the Gene Ontology (GO) annotation. The used GO biological process terms included “DNA binding activity” and “transcription regulator activity”. 1855 genes with both GO annotation were collected. Then, they were filtered for the ones with gene expression equal to 0 in any sample of RNA-seq data. Finally, 869 TFs were used for bi-clustering analysis.

We also performed TF binding site over-representation analysis for the target genes identified by bi-clustering analysis. Here, TF binding annotation were collected from RcisTarget package [30]. 490 out of 869 TFs were selected for evaluation by filtering the ones without binding motif annotation and the ones with less than 50 target genes. Then, the TFs identified by bi-clustering analysis were checked for motif enrichment. To evaluate the random occurrence of enriched TF binding sites, we randomly selected *n* brain-expressed genes, where *n* was equal to the number of TF-regulated target genes, and found the chance for false recovery was about 6% on average.

### 4.6 Bi-clustering analysis

We developed a bi-clustering algorithm to study patient divergence among AD patients. The philosophy behind this algorithm was to cluster the AD patients into subsets with different TF regulation statuses. To measure the regulation status, we selected a set of biomarker genes to indicate the TF activity, including both average co-expression correlation and number of TF-regulated genes. To make sure of confidence, we choose a set of strict parameters setting to define the dominant regulation, including the minimum size of TF-regulated patients *n* > 50, the minimum number of TF-regulated genes *m* > 30 and the minimum co-expression correlation *r* > 0.8. Only the result satisfying all these thresholds would be reported. As the process of bi-clustering analysis would generate a continuous number of patients and genes, the solution of bi-clustering analysis was reported under three scenarios: the solution with the maximum number of genes, the solution with the maximum number of patients and the solution with the maximum multiplied product values of patient and gene number. We evaluated them for their clinical association and found that the first one had overall better clinical consistency, which better indicated the status of regulation loss.

The full description of the algorithm is available in Supplementary Methods. The R and C++ source for bi-clustering analysis is available: https://github.com/menggf/bireg.

### 4.7 regulatory degeneration burden

We used regulatory degeneration burden (RDB) to measure the degree of regulatory degeneration in a subject. Its calculation was based on the prediction result of bi-clustering analysis. In this process, each subject was denoted by a binary vector *x*_*i*_, in which 1 indicated transcription regulation of a corresponding TF and 0 indicated no regulation. RDB is defined as

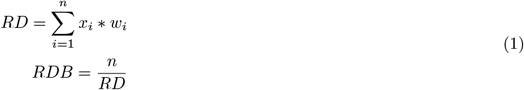

where, *n* is the number of TFs and *w*_*i*_ is a constant value that is only determined by regulation types. In this work, we tried different *w*_*i*_ values for WR TFs, ranging from 0 to 1, and evaluated them by fitting the clinical status of subjects. The evaluation results indicated the values from 0.5 to 1 were acceptable in most of the cases. In this work, we used the following setting:

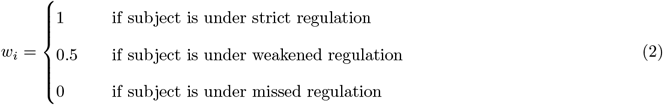

### 4.8 Peak calling

Paired-end reads of ChIP-seq and ATAC-seq DNA fragments were generated by Illumina Novaseq 6000 platform. On average, 20 million reads were available for each sample. The raw fastq files were firstly checked by *fastqc* and found overall good quality of the sequencing data. We used NGmerge to remove the adaptors [56]. Then, reads were aligned to human genome hg38 using bowtie2 [57]. The output sam files were filtered for reads with PCR duplicates, unmapped tags, non-uniquely mapped tags or with mismatches greater than 2. SAM files were then transformed into sorted and indexed bam files by samtools [58]. We applied macs2 [59] to identify the peaks under a cutoff of *q* < 0.05. For ATAC-seq data, we used a set of option “–broad –nomodel –shift 37 –extsize 73 –keep-dup all”.

For both AD and control group, we observed that the number of reported peaks are quite diverse among subjects. To identify the peaks with more confidence, we did cross-validation to select the peaks observed in more than one sample using “bedtools”. Each “*.bed” file was checked with other peak files using “bedtools intersect -wa -f -r 0.6”.

### 4.9 Differential peak analysis

Besides of peak calling analysis for each sample, we also did differential peak analysis to study the overall histone modification/open chromatin accessibility/TF binding intensity in AD and control samples. In this process, we applied an analysis pipeline proposed by S.J. Marzi et.al [20]. In peak calling steps, bam files of both AD and normal subjects were merged to maximize the power of peak calling. Under a parameter setting mentioned above, MACS2 [59] was used. Considering the fact that the histone modification marks usually spread broad regions, we set “–broad” option for the ChIP-seq data of H3K27ac, H4K16ac and ATAC-seq data. We validated and filtered the identified peaks by checking the peak overlap using the ones identified in the NIH RoadMap Epigenomics Consortium in all brain regions, including angular gyrus, anterior caudate, cingulate gyrus, BI.middle hippocampus, inferior temporal lobe, middle frontal lobe and substantia nigra. Next, we estimated the read abundance of peaks using the tools in `Rsubread` package. After filtering the peaks with read count of 0 or total reads number is less than 100, we did differential peak analysis to find the genomic regions with and gain. In this step, we used edgeR [55] to identify the peaks with altered TF binding affinity or open chromatin accessibility status. Different from S.J. Marzi et.al’s analysis pipeline, we consider the facts that the peaks loss and gain are not equal in AD samples and one-step differential peak analysis may cause under-estimation to the regulation loss. A better solution is to normalize the count data using only peaks without regulation loss or gain. Therefore, we applied a multiple-round analysis. In each round, we perform differential peak analysis and remove the differential peaks. This process was repeated until no new differential peak was reported anymore. The library size and dispersion information of count data were updated using the count matrix reported in the last round of differential peak analysis. Considering that the selected histone modification markers, open chromatin accessibility and TF binding sites were all associated with active regulation, we reported the peaks with increased read abundance as regulation gain and the peaks with decreased abundance as regulation loss at a cutoff of FDR < 0.05. Using `ChIPseeker` package [60], we annotated the differential peaks for their genome location and nearby genes in a arrange from -3000 bp to 1000 bp round transcription start sites.

### 4.10 Functional annotation

Our evaluation suggested that the regulators took dominant regulatory roles to their regulated genes. The functional involvement of regulators was inferred by annotation to the downstream-regulated genes. The R package `clusterProfiler` was implemented for enrichment analysis using Gene Ontology biological process terms. Under a cutoff of Benjamini *p* < 0.05, the enriched terms were selected and calculated for their frequency among all the regulators. After manually filtering the terms with overlapped functional annotation, terms with both significant enrichment and high frequency were identified to represent the functional involvement of regulators. The used terms were clustering using the hierarchical clustering method based on the *p*-values reported by enrichment analysis. To smooth the heatmap visualization, the *p*-values were transformed with −*log*10. If the −*log*10(*p*) value is greater than 6, they were set to 6. We also did the functional annotation for each regulator by text-mining tools, including Ingenuity Pathway analysis (IPA), GeneCards (https://www.genecards.org/) and PumMed, to evaluate the functional consistency among predictions from different sources.

### 4.11 Enrichment analysis

Enrichment analysis was performed to evaluate the statistical significance of feature overlaps, such as genes associated with the same biological process. For *k* input genes, the number of genes with certain feature is *x*. Among *n* whole genomic genes, the number of genes with such feature is *p*. Fisher’s exact test can evaluate if the observed *x* genes resulted from random occurrences. We used the R codes to calculate statistical significance:

> *m* = *matrix*(*c*(*x, k* − *x, p* − *x, n* − *k*), *ncol* = 2, *byrow* = *T*)

> *p* = *fisher*.*test*(*m, alternative* = “*greater*”)$*p*.*value*

### 4.12 Co-expression network analysis

We used WGCNA, as the implementation of co-expression network analysis, to predict gene modules of expression data by following the protocol introduced by the official document in https://labs.genetics.ucla.edu/horvath/htdocs/CoexpressionNetwork/Rpackages/WGCNA/Tutorials/index.html. Considering the fact that WGCNA has some parameter settings while no golden standard to optimize them, we test different setting one by one and chose the setting combination where the best clinical association was observed as the final parameter setting. The summarized profiles or eigengenes of modules were investigated for their clinical association by Spearman’s correlation. Except for the gray module, modules with the best clinical correlation were selected for evaluation.

### 4.13 Cellular deconvolution and the association with RDB

The bulk RNA-seq data used in this study were analyzed for in silico deconvolution to estimate the cellular compositions. In this process, R package *granulator* was used for analysis. We collected the reference profile of 6 brain cell types from published single-cell RNA-seq study to AD [61]. The measured cell type included oli (oligodendrocytes), mic (microglia), ast (astrocytes), opc (oligodendrocyte progenitors), in (inhibitory neurons), ex (excitatory neurons). The used marker genes included SYT1, SNAP25,GRIN1, SLC17A7, CAMK2A, NRGN, GAD1, GAD2, MBP, MOBP, PLP1, PDGFRA, VCAN, CSPG4, CD74, CSF1R, C3, AQP4, GFAP, FLT1, CLDN5. The calculated cellular proportions were evaluated for their partial correlation with RDB. Using the AD-related clinical traits as covariates, the partial correlations were calculated with pcor function in *ppcor* package. Additionally, we also calculated the partial correlations between RDB and clinical traits by controlling the cellular composition.

mathys2019single

### 4.14 Experimental Materials for HDAC1 studies

Antibodies of HDAC1(D5C6U), APP (E8B3O), *p*-Tau^*Ser*^ (D7U2P), *p*-^*TauThr*205^ (E7D3E), *p*-Tau^*Thr*181^ (D9F4G) and *p*-Tau^*Ser*181^ (D2Z4G) were purchased from Cell Signaling Technology (Danvers, MA, USA). Antibodies of GAPDH and Tau were purchased from Abmart (Shanghai, China). Human Aβ_1−42_ (Amyloid Beta1-42) and Human Aβ_1−42_ (Amyloid Beta1-40) ELISA kit was obtained from ebioscience (Sangon Biotech, Shanghai, China) HDAC1 plasmid were acquired from Beyotime (Haimen, China). Lipofectamine 3000 was purchased from Invitrogen (Carlsbad, CA, USA). Small interfering RNA (siRNA) targeting HDAC1 was purchased from GenePharma (Shanghai, China).

### 4.15 Plasmids transfection and RNA interference

The cell lines of HEK/APPsw cells were obtained from Cell Bank of Shanghai institute of Cell biology, Chinese Academy of Sciences (SIBS, CAS). The cell was cultured in DMEM medium with 10% fetal bovine serum and 1% penicillin/streptomycin with 5% CO2 at 37 ∘C. HEK/APPsw cells (2×106 cells) were plated in 6-well plates per well (Corning). After cultured overnight, the Flag-HDAC1 plasmid was transfected into HEK/APPsw cells with Lipofectamine 3000 according to the manufacturer’s protocol. For siRNA-mediated silencing, cells were transfected with 100 nmol/L of target siRNA and a control siRNA using Lipofectamine 3000. 48 hours post-transfection, the protein expression was analyzed by immunoblotting and ELISA. siRNAs used in this study included:

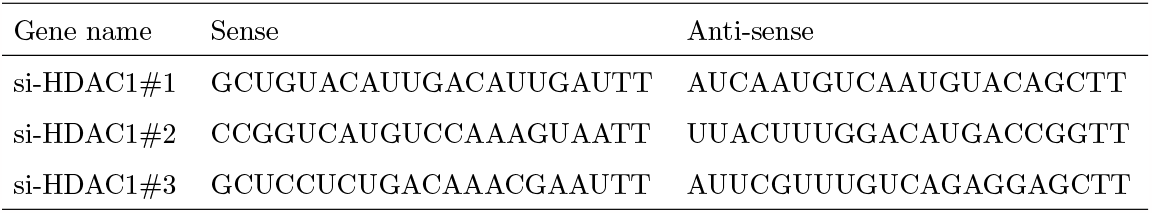

### 4.16 Protein Extraction, Western Blot Analysis and ELISA

Total cell protein was extracted using RIPA Lysis Buffer supplemented with 1x protease inhibitor and 1x phosphatase inhibitor. Protein was loaded for gel electrophoresis and transferred onto polyvinylidene difluoride membranes. The membranes were probed with the corresponding primary antibodies at 4∘C overnight and horseradish peroxidaseconjugated secondary antibodies at room temperature for 1 h. Detection was performed with an Odyssey infrared imaging system (LiCor Biotechnology, Lincoln, NE). The levels of Aβ_1−42_ and Aβ_1−40_ in cells were detected by ELISA kit according to the manufacturer’s protocol.

### 4.17 RNA isolation and quantitative real-time PCR

Total RNA was isolated by RNAiso Plus (Takara, Dalian, China). RNA (1μg) was reverse transcribed using the Prime Script RT reagent Kit (Takara, Dalian, China). Quantitative RT-PCR was performed using Roche LightCycler480 and the sequences of the primers are indicated in Table 2. The relative levels of assayed mRNAs were calculated with the comparative CT method using GAPDH expressions as endogenous control and were normalized to the non-treated control. Primer sequences used in this study included:

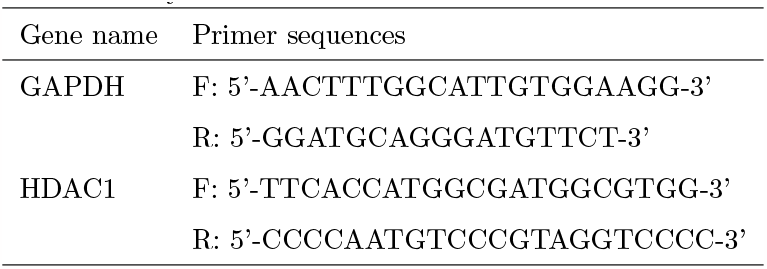

### 4.18 HDAC1 over-expressed transgenic mice

CAG-LSL-HDAC1-3FLAG-PolyA knockin mice were generated by GemPharmatech using the CRISPR–cas9 system. C57BL/6J and CAG-LSL-HDAC1-3FLAG-PolyA knock-in mice were purchased from GemPharmatech(Nanjing, China), maintained uniformly by the Shanghai Laboratory Animal Center (Shanghai, China) of the Shanghai University of Traditional Chinese Medicine (SHUTCM), and housed under specific pathogen-free conditions. Animals were housed in an environment-controlled room (temperature 20-25∘C, relative humidity 55-65%, and 12 h light / 12 h dark cycle) with free access to water and feed. All animal experiments were in accordance with international ethical guidelines and National Institutes of Health Guidelines on the Care and Use of Laboratory Animals, which were approved by the Animal Experimental Ethics Committee of the Shanghai University of Traditional Chinese Medicine (SHUTCM). We made every effort to minimize any pain to the animals.

### 4.19 Mouse behaviour test

The 10-month-old mice were evaluated in NOR and MWM tests. Before the behaviour tests, mice were acclimatized to the behavioural testing room for 2 h. All experiments were completed between 8:00 and 18:00, and the behavioural performance of the mice was recorded and analyzed by an automatic tracking system (EthoVision-XT; Noldus, Wageningen, Netherlands). The test box was sequentially cleaned with 70% and 30% ethanol before the next mouse was introduced to the box. The spatial learning and memory abilities of the mice were assessed through the MWM task. The device is a circular white pool (120 cm diameter x 50 cm depth) filled with water dyed white with TiO_2_, and with temperature maintained at 22 ∘C. A 10-cm-diameter platform was placed 1 cm below the water surface at a fixed position. Mice were trained with 3 trials per day for 5 consecutive days. Each trial lasted 60 s or until the mouse found the platform. If the mouse did not find the platform during the allocated time period, the experimenter directed the mouse to the platform. After each trial, the mouse was placed on the platform for 15 s.On the 6th day, the platform was removed from the pool, and each mouse was allowed to swim freely for 60 seconds. The amount of time spent in the quadrant of the original platform location was calculated as an index of long-term spatial memory. Statistical analysis was conducted by using GraphPad Prism 8.0 software (La Jolla, CA, USA). Data were presented as means *±* Standard Error of Mean(SEM) from at least three-independent experiments. All experiment data were statistically evaluated by using t-test or one-way ANOVA with Tukey’s multiple comparison tests.

### 4.20 Brain immunohistochemistry

Mice (n = 3 per group) were deeply anaesthetized with 10% Chloral hydrate (0.25 g/kg body weight) and then immediately perfused with normal saline followed by 4% paraformaldehyde (PFA) in 0.1 mol/L phosphate buffer. Harvested brain tissues were postfixed in the same solution at 4 ∘C for 4 hours and then dehydrated in a graded series of sucrose solutions until they were fully permeated. Frozen brain sections were cut 30-μm thick in a vertical plane using a cryostat microtome (CMS3050S, Leica Microsystems, Nussloch, Germany), and IHC staining was performed on 3 sections per mouse. Slices were incubated with the primary antibodies of Aβ overnight at room temperature and the appropriate biotinylated secondary antibodies (1:200, Vector Laboratories, Burlingame, CA, USA) for 1 hour at room temperature, followed by incubation with an avidinbiotin-peroxidase complex (ABC kit; Vector Laboratories, Burlingame, CA, USA). After rinsing with PBS (10 minutes three changes), the sections were incubated with a diaminobenzidine (DAB) kit (Vector Laboratories, Burlingame, CA, USA). Slices were stained and observed using a light microscope (Leica, Wetzlar, Germany). Images were obtained using ImageJ and the threshold intensity was manually set at a constant value for all images. Pixel counts of Aβ load were derived from the average of three adjacent sections per animal.

## 5 Data Available

The ATAC-seq and ChIP-seq data were publicly available in the Gene Expression Omnibus (GEO) database with the ID of GSE129041. The RNA-seq data were available with the ID of GSE189725. The Perl and R source code for differential peak analysis is available:https://github.com/menggf/bireg/tree/master/code.

## 6 Acknowledgements

We are grateful to the China Brain Bank of Zhejiang University School of Medicine and Chinese Brain Bank Center in Wuhan for providing human brain material. The results published here are in part based on data obtained from the AMP-AD Knowledge Portal (doi:10.7303/syn2580853). We appreciate their generous contribution to the studies of Alzhiemer’s disease. We thank Dr. Lichun Jiang, Dr. Ruping Sun for reading this manuscript and giving us many useful suggestions. We also thank Dr. Guhan Nagappan, Dr. Xiaoming Guan, DR. Bai Lu for valuable comments. The work was supported by NSFC (81973706, 81520108030, 21472238), Shanghai Engineering Research Center for the Preparation of Bioactive Natural Products (16DZ2280200), the Scientific Foundation of Shanghai China (13401900103, 13401900101), the National Key Research and Development Program of China (2019YFC1711000, 2017YFC1700200), Innovation Team and Talents Cultivation Program of National Administration of Traditional Chinese Medicine (No: ZYYCXTD-D-202004), Shanghai Frontiers Science Center of TCM Chemical Biology, Innovation Team and Talents Cultivation Program of National Administration of Traditional Chinese Medicine. (No: ZYYCXTD-D-202004)

## Notes

### Competing Interest Statement

The authors have declared no competing interest.

