## Supplementary material for "Computational divergence analysis reveals the existence of regulatory degeneration and supports HDAC1 as a potential drug target for Alzheimer’s disease": Supllementary Materials

### Regulatory degeneration contributes to development of Alzheimer's disease and supports epigenetic regulator HDAC1 as a potential drug target

Guofeng Meng et. al

#### Supplementary information

- Supplementary Methods
  - Chromatin Immunoprecipitation (ChIP-seq)
  - Assay for Transposase-Accessible Chromatin using sequencing (ATAC-seq)
  - Bi-clustering algorithm
- Supplementary Results
  - Ability to identify the subsets of patients under different regulation context
  - Evaluation to false discovery ratio of bi-cluster analysis
  - Impact of different cutoffs on the bi-clustering analysis result
  - Evaluation using brain normal tissues
- Supplementary figures
  - Figure S1: Clinical information of AD subjects;
  - Figure S2: Regulation status of predicted regulators;
  - Figure S3: WR regulators more associated with clinical outcomes than MR;
  - Figure S4: WR and MR regulator take dominant regulatory roles;
  - Figure S5: Ubiquitin-proteasome system is disturbed by transcriptional regulation loss;
  - Figure S6: RDB is associated with clinical outcome;
  - Figure S7: The correlation among different clinical traits and cell compositions.
  - Figure S8: RDB better indicates clinical outcomes than genes, transcription factors and WGCNA modules;
  - Figure S9: Genomic distribution of TF binding sites and ATAC-seq peaks
  - Figure S10: ChIP-seq peak loss of three TFs is overlapped with the predicted dysregulated genes

– Figure S11: Regulation loss indicated by histone modification mark, H4K16ac;

– Figure S12: Differential expression and clinical association results of 11 HDAC genes;

– Figure S13: The water mirror maze results for (a) the spent time and (b) swimming distance in the target quadrant.

### 1 Supplementary Methods

#### 1.1 Chromatin Immunoprecipitation (ChIP-seq)

The whole experiment was performed following the published protocols. Firstly, weigh 30-40 mg of frozen tissue in a petri dish. Chop tissue into small pieces (between 1-3 mm3 using a scalpel blade. Add 1 ml of ice-cold PBS with protease inhibitor cocktail and disaggregate the tissue using a dounce homogenizer (or pestle) to get a homogeneous suspension. Transfer the tissue suspension into a 1.5 ml tube and centrifuge at 1,300 rpm for 5 min at 4°C. Gently discard the supernatant and keep the pellet. Resuspend the pellet in 1 ml of ice-cold PBS containing formaldehyde to a final concentration of 1% (37% in stock) at room temperature. Rotate tube for 10-15 min at room temperature. Stop the cross-linking reaction by adding fresh glycine to a final concentration of 0.125 M. (2.5 M in stock). Continue to rotate at room temperature for 5-10 min. Centrifuge samples at low speed (1,300 rpm) at 4°C. Wash the pellet with 1 ml of ice-cold PBS (plus protease inhibitors). Aspirate the supernatant and resuspend the pellet in 1 ml of ice-cold PBS (plus protease inhibitors). Centrifuge at low speed (1,300 rpm) at 4°C and discard the supernatant. Repeat the washing one more time. Flash freeze the tissue pellet in liquid nitrogen and store at -80°C.

Chromatin was sonicated by Bioruptor (Diagenode, Belgium) under the condition as follows: 20 cycles of 30s on, 60s off. The sonicated chromatin was spun down at 12,000 rpm for 10 min at 4 °C to collect the chromatin. Add antibody (10 $\mu$g) to the soluble chromatin (80 $\mu$g), and incubate at 4 °C with rotation overnight. Protein A/G Dynabeads (10 $\mu$l of beads per 1 $\mu$g of antibody, Life Tech, 10004D) were washed three times with Low Salt Wash Buffer. The washed Dynabeads were added to the soluble chromatin and antibodies, incubated at 4 °C for 4 h with rotation. The magnetic Dynabeads were pelleted by placing the tubes in a magnetic rack and were sequentially washed once with Low Salt Wash Buffer, once with High-salt Wash Buffer, once with LiCl Wash Buffer. Wash the beads three times with TE buffer. Remove any supernatant remaining after the last washing. The beads were resuspended in 150 $\mu$l of Elution Buffer (50 mM Tris-Cl pH 8.0, 10 mM EDTA, 1% SDS), followed by incubation for 15 min at 65 °C. Repeat step 7 again, combine the elution, then you have 300 $\mu$l of the eluted DNA solution. Add 1 $\mu$l of high concentration RNase A (10 mg/ml) to the eluted DNA solution and the input samples (500 $\mu$l), respectively. Incubate them at 37 °C for 1 h. Reverse formaldehyde crosslinks by respectively adding 12.5 $\mu$l or 55 $\mu$l of 5 M NaCl to the eluted DNA solution and the Input samples to a final concentration of 0.2 M. Incubate samples at 65 °C (650 rpm) for more than 8 h (less than 16 h). Add both 80.5 $\mu$l of ddH2O and 5 $\mu$l of 20 mg/ml Proteinase K to the eluted DNA solution, and just 5 $\mu$l of Proteinase K to the Inputs. Incubate at 56 °C for 2 h. Extract once with 400 $\mu$l of phenol/chloroform/isoamyl alcohol, and once with 400 $\mu$l of chloroform/isoamyl alcohol (optional). Transfer 355 $\mu$l of supernatant to a new tube. Add 55 $\mu$l or 40 $\mu$l of 3 M NaAc (pH 5.2, 0.3 M final) to the Input samples and eluted DNA solution, 5 $\mu$l of glycogen each sample and mix well. Add 800 $\mu$l (two-fold volume) of absolute ethanol. Precipitate at -20 °C overnight or at -80 °C for 4 h. Centrifuge at 14000 rmp for 20 min at 4 °C. Wash once with 75% ethanol and store at -20 °C.

The used the antibody includes (1) ATF4, CTS,11815S (lot#4); (2) Olig2, RnD, AF2418 (lot#UPA0718031); (3) THRA, SantaCruz, SC-56873 ((lot#J1614).

#### 1.2 Assay for Transposase-Accessible Chromatin using sequencing (ATAC-seq)

ATAC-seq was performed in GENEWIZ company following the protocol introduced in [1, 2]. Place frozen tissue into a pre-chilled 2 ml Dounce with 2 ml cold nuclei lysis buffer. Allow frozen tissue to thaw for 5 minutes. Dounce with A pestle until resistance goes away ( 10 strokes). Dounce with B pestle for 20 strokes. Pre-clear larger chunks by pelleting at 100 RCF for 1 min in a pre-chilled centrifuge.Count nuclei using Trypan blue staining and aliquot nuclei for ATAC reaction. Harvest and count cells (counting protocol to be defined by the user; see APPENDIX3F). Cells should be intact and in a homogenous, single-cell suspension; Centrifuge 50,000 cells 5 min at 500 x g, 4°C. The number of cells at this step is crucial, as the transposase-to-cell ratio determines the distribution of DNA fragments generated. Remove and discard supernatant. Wash cells once with 50 $\mu$l of cold PBS buffer. Centrifuge 5 min at 500x g, 4°C. Remove and discard supernatant. Gently pipet up and down to resuspend the cell pellet in 50 $\mu$l of cold lysis buffer. Centrifuge immediately for 10 min at 500 x g, 44°C. Discard the supernatant, and immediately continue to transposition reaction. Make sure the cell pellet is set on ice. To make the transposition reaction mix, combine the following: TD (2x reaction buffer from Nextera kit) 25 $\mu$l; TDE1 (Nextera Tn5 Transposase from Nextera kit) 2.5 $\mu$l; Nuclease-free H2O 22.5 $\mu$l. Resuspend nuclei pellet (from step 5) in the transposition reaction mix. Incubate the transposition reaction at 37°C for 30 min. Gentle mixing may increase fragment yield. Immediately following transposition, purify using a Qiagen MinElute PCR Purification Kit. Elute transposed DNA in 10 $\mu$l Elution Buffer (Buffer EB from the MinElute kit consisting of 10 mM Tris·Cl, pH 8). To amplify transposed DNA fragments, combine the transposed DNA (10 $\mu$l), nuclease-free H2O (10 $\mu$l),25 $\mu$M PCR Primer 1 (2.5 $\mu$l), 25 $\mu$M Barcoded PCR Primer 2 (2.5 $\mu$l), NEBNext High-Fidelity 2x PCR Master Mix (2 5 $\mu$l). Thermal cycle as follows 72°C,5 min, 1 cycle; 98°C,30 sec; 98°C, 10 sec 5 cycles; 63°C, 30 sec; 2°C, 2 min; 4°C.

#### 1.3 Bi-clustering algorithm

In this work, we developed a bi-clustering algorithm to study AD patient diversity in TF-mediated transcription regulation. The philosophy behind this algorithm is to find the patient subset with different transcription regulation context. Let's demonstrate its necessity with following R codes:

```r
> x1= rnorm(100,1,1) # a simulated expression vector for 100 subjects as population 1;
> y1= x1 + rnorm(100,1,0.5) # to generated a vector of regulators
> cor(x1,y1)
# output 0.8529, indicated strong regulation

> x2= rnorm(50,1,1) # second simulated vector of 50 subjects as population 2;
> y2= x2 + rnorm(50,1,3) # to generated a vector of regulators
> cor(x2,y2)
# output 0.3015, indicated weak regulation

> x=c(x1, x2) # merge population 1 and 2
> y=c(y1, y2) # merge regulators
> cor(x,y)
```

### output 0.4648, indicated a mediate regulation in mixed population

Above demo indicates that patient mixture can conceal the true TF-gene regulation. There should be tools to study cohort structure based on their regulation status.

To do it, we make two assumptions: (1) we can find a set of marker genes to indicate the TF regulation activity; (2) AD patients can be clustered into subsets under different regulation context. Considering the fact that bi-clustering is an NP (nondeterministic polynomial time) problem, we designed an approximating implement. It always starts with a TF gene ($g_i$) and a gene expression matrix. The patient subsets are identified by following steps:

1. The co-expression correlation, measured by Spearman's method was calculated for TF-gene pairs, $r_{i,j} = cor(e_{g_i}, e_{g_j})$, where $e_{g_i}$ was the expression vector for TF gene $g_i$ and $e_{g_j}$ was the expression vector for gene $j$;
2. The genes were ranked by $r_{i,j}$ and the top $m^{'}$ genes were selected as the initial gene set $G = \{g_1, g_2..g_{m'}\}$;
3. For each gene $g_j$ in $G$, the subjects were ranked based on the degree of correlation improvement after subject removal, where the correlation improvement was described by expression rank difference: $d_{i,j} = |d_i - d_j|$;
4. A voting step was used to select the subject with the best Spearman's correlation improvement: $argmin\{\sum_{j=1}^{m} d_{i,j}\}$. In this process, we did following steps:
    (a) To calculate the rank differences between TF and genes, and assigned the priority rank to remove in next round;
    (b) To count the rank occurrence in the windows of 50;
    (c) To selected the one with least value of rank occurrences as subject to remove;
5. Remove the selected subject and recalculate the Spearman's correlation;
6. Repeat above process until more than $m$ genes in more than $n$ subjects with co-expression correlation greater than a predefined cutoff $r_{min}$.

This process is repeated for all the TF genes. To make sure of the confidence of bi-clustering analysis results, we chose a set of strict parameters setting for our data based on evaluation analysis, including $n > 50$, $m > 30$ and $r_{min} = 0.8$. Only the result satisfying these thresholds would be reported. As the process of bi-clustering analysis would generate continuous number of patient and gene number combination, we checked the output results under three scenario: the solution with the maximum number of genes, the solution with the maximum subjects and the solution with the maximum product values of patient and gene number. We evaluated them using clinical association and found that to maximize the number of genes had overall better clinical consistency, which might better indicate the status of regulation loss. Therefore, in this work, we chose the gene-subject combination when maximum gene number as the solution of bi-clustering analysis.

#### 2 Supplementary Results

We evaluated the performance of our bi-clustering algorithm using both simulated data and real brain data mentioned in this manuscript. In this step, we performed four evaluations, including (1) ability to identify the subsets of patients under different regulation context; (2) evaluation to false discovery ratio of bi-cluster analysis; (3) impact of co-expression correlation cutoffs on analysis results; (4) evaluation using independent normal brain tissues. Additionally, we also did one evaluation to the existence of neuronal loss in the data used in this work.

#### 2.1 Ability to identify the subsets of patients under different regulation context

To test if our bi-clustering algorithm grouped the subset of patients under different regulation context, we generated a simulated data under different regulation context with r codes are like following:

```
> x1=rnorm(200,1,1)
> mx11 = t(sapply(1:100, function(i) x1 + rnorm(200,1,0.4 + rnorm(1, 0, 0.01))))
> mx12 = t(sapply(1:100, function(i) x1 + rnorm(200,1,1.5 + rnorm(1, 0, 0.01))))
> mx21 = t(sapply(1:14900, function(i) x1 + rnorm(200,1,5 + rnorm(1, 0, 0.01))))
> mx22 = t(sapply(1:14900, function(i) x1 + rnorm(200,1,5 + rnorm(1, 0, 0.01))))
> mx=rbind(cbind(mx11, mx12), cbind(mx21, mx22))
> # the matrix has a form of
> # mx11 | mx12
> # ——    ——
> # mx21 | mx22
> row.names(mx) <- paste("g",1:15000,sep="")
> colnames(mx) <- paste("p",1:400,sep="")
```

In this simulated data, "mx11" represents the subset of gene-patient combination under strict regulation (median $r = 0.86$) while "mx12" represents the subset with regulation loss (median $r = 0.31$). "mx21" and "mx22" are the gene without regulation and they are used as background. We generated such simulated data for 100 times and performed bi-clustering analysis. We summarized the analysis results and found that our algorithm accurately predicted all the true clusters for both patient and genes (100% accuracy, no figure is showed).

As a technical evaluation, this result suggested that our bi-clustering algorithm has good ability to identify subjects under different regulation contexts and our algorithm can identify the subset of AD patients under different regulation context.

#### 2.2 Evaluation to false discovery ratio of bi-cluster analysis

Bi-clustering analysis is a NP (nondeterministic polynomial time) problem in algorithm design. Our tool applies an approximating implement by moving out subjects one by one to reach a maximum number of TF-regulated genes (or TF-gene pairs) under a strict threshold. One concern is about its false discovery ratio or if its prediction results are only due to technical bias. We performed `two` evaluations using simulated data and real data.

In the first evaluation, we randomly shuffled the gene expression data so that each gene had true gene expression values but wrong assignment to patients. In such simulated data, the TF-gene regulation was supposed to be completely disrupted by shuffling. We did bi-clustering analysis to check if any TF-gene regulation could be falsely identified by our algorithm. We relaxed the co-expression correlation cutoff to to $|r| > 0.4$ and the others are the exact same

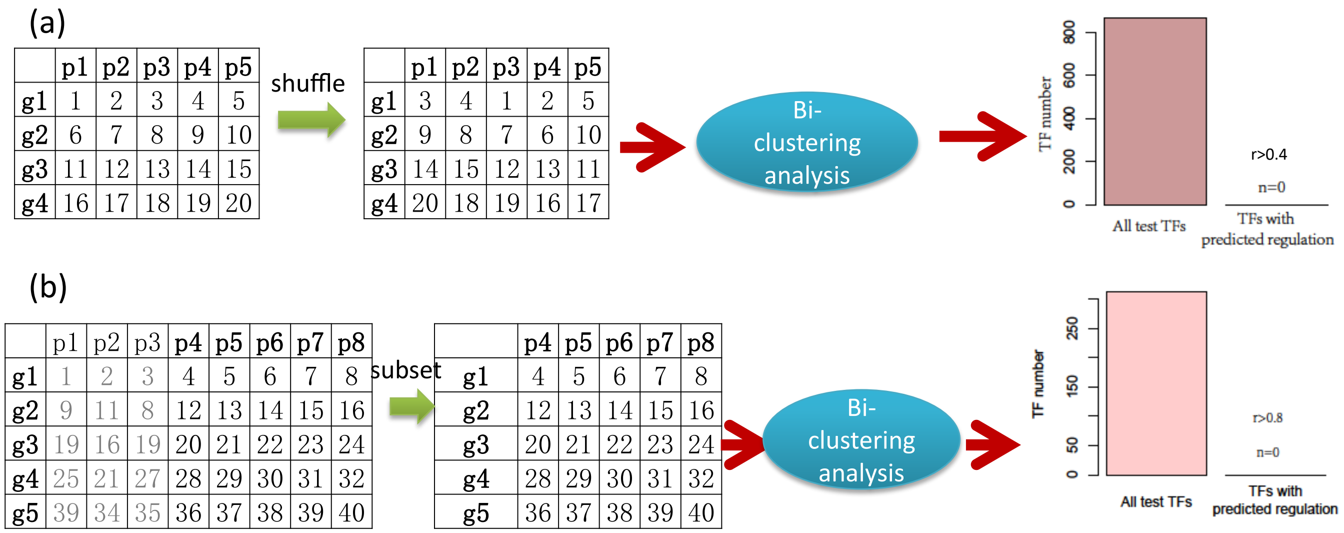


Figure 1: Evaluation to false discovery ratio of predicted regulation. (a) The gene expression data were shuffled to disrupt the TF-gene regulation and then checked if any false regulation would be predicted. Here, we did not observed any false prediction even when loosed cutoff was used. (b) We filtered the subjects with TF regulation (e.g. p1,p2,p3) and did another round of bi-clustering analysis using the only non-regulated subjects. We did not observed any new TF-gene regulation.

parameter setting mentioned in the manuscript. The whole evaluation process, from data shuffling to bi-clustering analysis, was repeated for 100 times. We summarized the results and did not find any predicted TF-gene regulation, including weakened and missed regulation (see Figure 1(a)). This result suggests that the predicted regulation loss is not due to technical bias of strict cutoffs.

We did another evaluation by performing second-round bi-clustering analysis to the subjects without TF regulation. The TF regulation statuses were determined based on the bi-clustering analysis results using AD expression data. We extracted subjects with weakened or missed TF regulation to form a new dataset as the input. To make sure of good confidence, we filtered the datasets with less than 100 subjects. Using the same parameter setting, including a cutoff of $|r| > 0.8$, we did second round of bi-clustering analysis to check if there was any new WR or MR regulation patterns. This process was performed for four brain regions, including 316 TFs in total. We did not find any pattern of WR or MR for studied TFs, where some samples were under strict TF-gene regulation while regulation loss was observed in remaining samples (see Figure 1(b)). This result suggests that there is no alternative solution for the same TFs.

Overall, our evaluation suggests that the predicted TF-gene regulation is almost impossibly resulted from technical biases and there is no alternative solution for bi-clustering analysis results.

#### 2.3 Impact of different cutoffs on the bi-clustering analysis result

In bi-clustering analysis, we have to set some arbitrary cutoffs to predict the TF-gene regulation. Among them, co-expression correlation is the most critical one. It needs an evaluation for the cutoff selection.

Using the AD expression data mentioned in the manuscript, we did bi-clustering analysis at four cutoffs, including $|r| > 0.6, 0.7, 0.8, 0.9$. We summarized the types of predicted TF-gene regulation (see Figure 2(a)). At a cutoff of $|r| > 0.6$, 40% of TFs were predicted with dominant regulation types. On the contrary, 50% of TFs take no regulatory roles at a cutoff $|r| > 0.9$. This result suggests that cutoff selection has great influence on the analysis results. Although

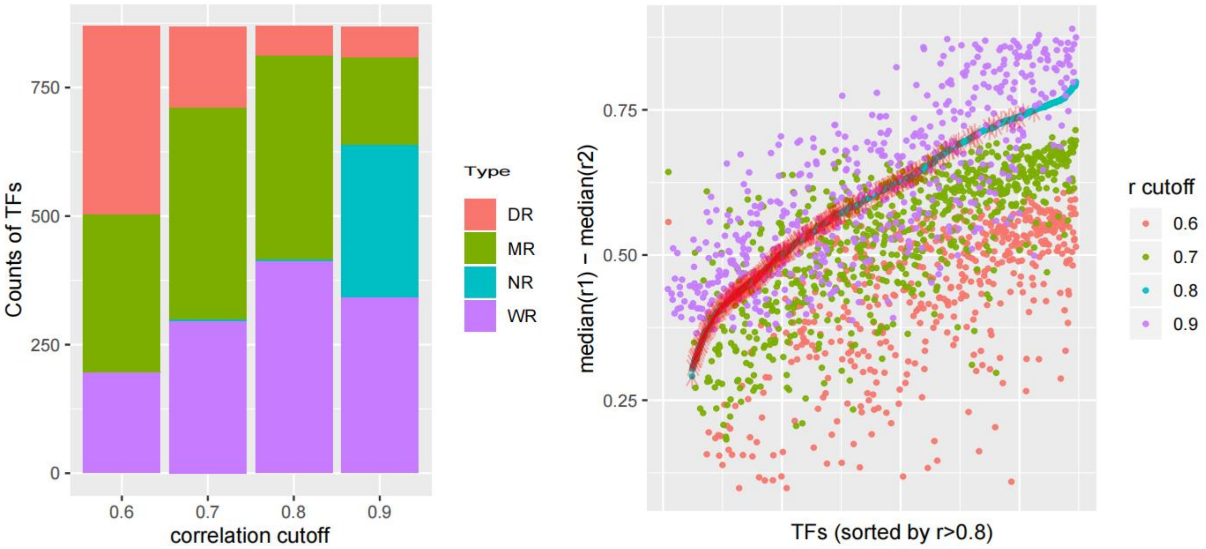


(a)

(b)

Figure 2: Impact of different cutoffs on the bi-clustering results. (a) The frequency of TF regulation types predicted at four different cutoffs. We observed that when $|r| > 0.6$ or $|r| > 0.9$ was set, about half of TFs were assigned with types of DR or NR, and they would not be evaluated for weakened or missed regulation, leading to low power of bi-clustering analysis. (b) Weakened and missed regulator predicted at 4 cutoffs. The y-axis indicated the co-expression correlation difference between TF-regulated subjects (r1) and the ones with TF regulation loss (r2). Red arrows mark the DR TFs predicted at a cutoff of $|r| > 0.6$. We found that most of DR regulation predicted at $|r| > 0.6$ could get weakened or missed at $|r| > 0.8$, which suggested the missed power at $|r| > 0.6$.

there is no golden standard to choose the proper cutoff, it seems that that too strict or loosed cutoff would result to more false negative discovery. Under our setting, only MR and WR TFs are evaluated for regulation loss. Too loosed or strict cutoffs could lead to low power to identify AD relevant TF regulation.

To further evaluate the cutoff selection, we checked if the DR TFs predicted at $|r| > 0.6$ had regulation loss at stricter cutoffs. In Figure 2(b), we show their regulation status at a cutoff of $|r| > 0.8$. We found that most of DR TFs predicted at $|r| > 0.6$ could get lost or weakened at $|r| > 0.8$ and the correlation value differences are big enough for a conclusion of regulation loss. This result suggests that the prediction at $|r| > 0.6$ has many false negative discoveries. We also evaluated the cutoff of $|r| > 0.7$ and still found more false DR regulation, which caused the lower power than the prediction at $|r| > 0.8$. On the contrary, many regulation loss were assigned with types of NR at $|r| > 0.9$, suggesting a the missed power in identifying regulation loss.

Overall, our results suggest too loosed or strict cutoffs can lead to low power; $r > 0.8$ is a good cutoff for the data used in this study.

#### 2.4 Evaluation using brain normal tissues

We test if the prediction of our algorithm could reflect the disease regulation status. We evaluated it based on an assumption that there should be less regulation loss prediction in normal brain tissues than AD samples. We collected RNA-seq data of three normal brain tissues from GTEx project (`https://gtexportal.org/home/`), include the regions of frontal cortex, cerebellum, cerebellar hemisphere. We did the bi-clustering analysis using our algorithm under the same parameter setting. In Figure 3(a), we showed the frequency distribution of predicted regulation types. We found

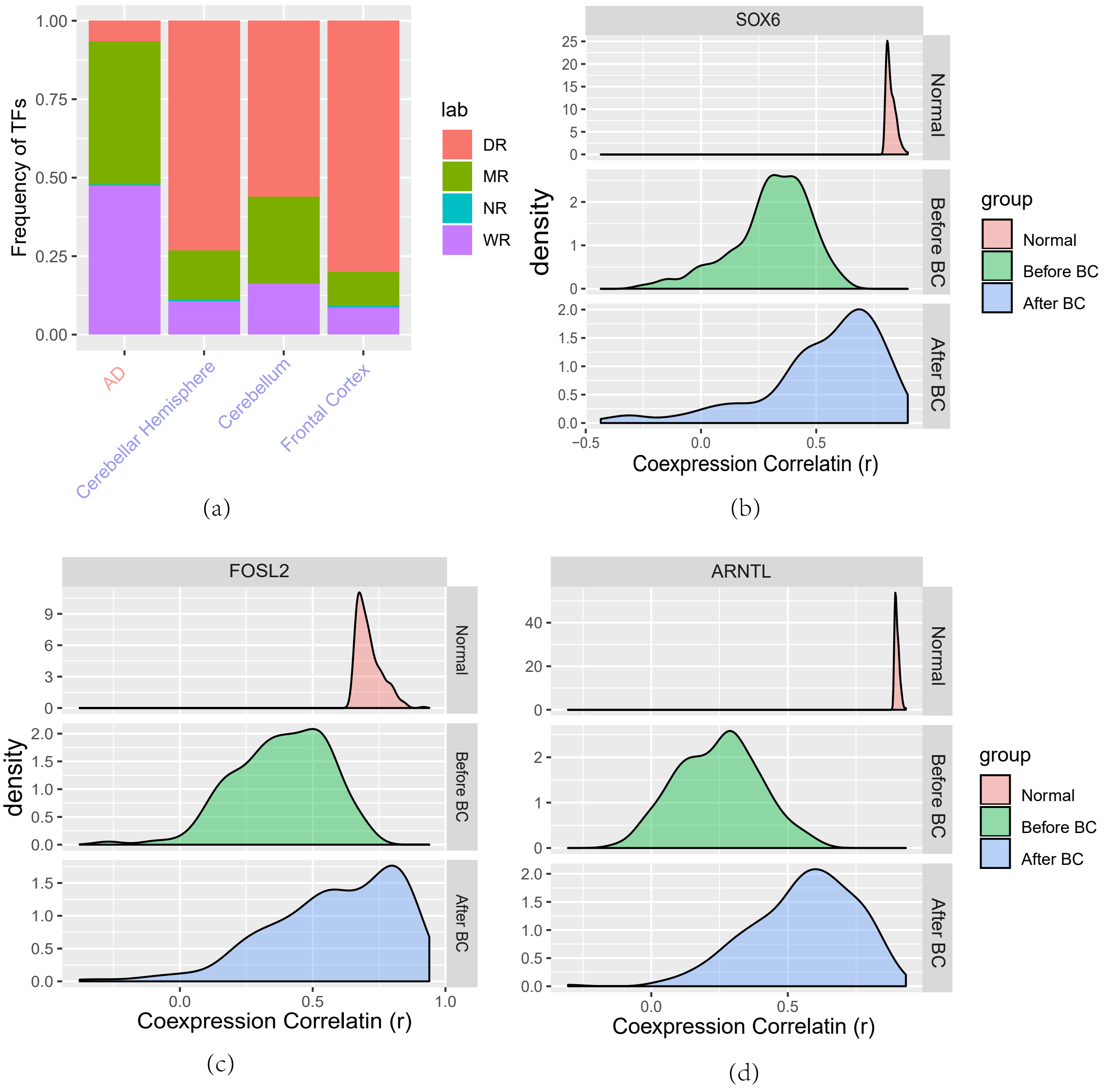


Figure 3: Bi-clustering analysis can recover the TF gene regulation in the normal brain tissue. (a) Bi-clustering analysis reports more DR regulation in all three normal brain tissues than the AD sample. This result suggests that regulation loss is less observed among normal cohorts. (b-c) Co-expression correlations of TF-gene pairs were improved after bi-clustering analysis. TF-gene regulation were firstly predicted by co-expression analysis to normal brain tissues and then their correlations were checked before and after bi-clustering analysis to AD samples. Here, three exemplary TFs were showed and we observed clear improvement of overall co-expression profiles after bi-clustering analysis. This result suggests that bi-clustering analysis could recover the TF-gene regulation in the normal brain tissues.

that DR was major regulation type of normal brain regions, account for 50%-70% of expressed TFs. The observed regulation loss was more likely due to the non-disease relevant factors, e.g. cell types and ages. Comparing to the results of AD data, this result is consistent with our assumption about less regulation loss.

Another evaluation was to check if bi-clustering analysis could recover the regulation discovered in the normal tissues. This evaluation is based an assumption that AD is associated with disruption of TF-gene regulation of normal brain. Therefore, we firstly collected the TF-gene regulation of normal brain tissues by co-expression analysis to normal brain expression data. The top 200 positively correlated genes were selected as biomarkers for TF regulatory activity in normal brain. Then, we evaluated the biological relevance of bi-clustering analysis results by checking if the normal TF-gene regulation could be recovered by our algorithm. Therefore, we compared their TF-gene co-expression correlation changes before and after analysis. Figure 3(b-d) showed three exemplary TFs and their correlation distribution with the biomarker genes. We observed that the co-expression correlation was greatly improved after bi-clustering analysis. These results suggest that the bi-clustering analysis can recover the true TF mediated regulation from the AD patients; the predicted regulation loss is relevant to disease.

Overall, our evaluations suggested that regulation loss was more observed in AD patients; our algorithm can recover the regulation of normal brain tissues from AD samples.

### 3 Supplementary figures

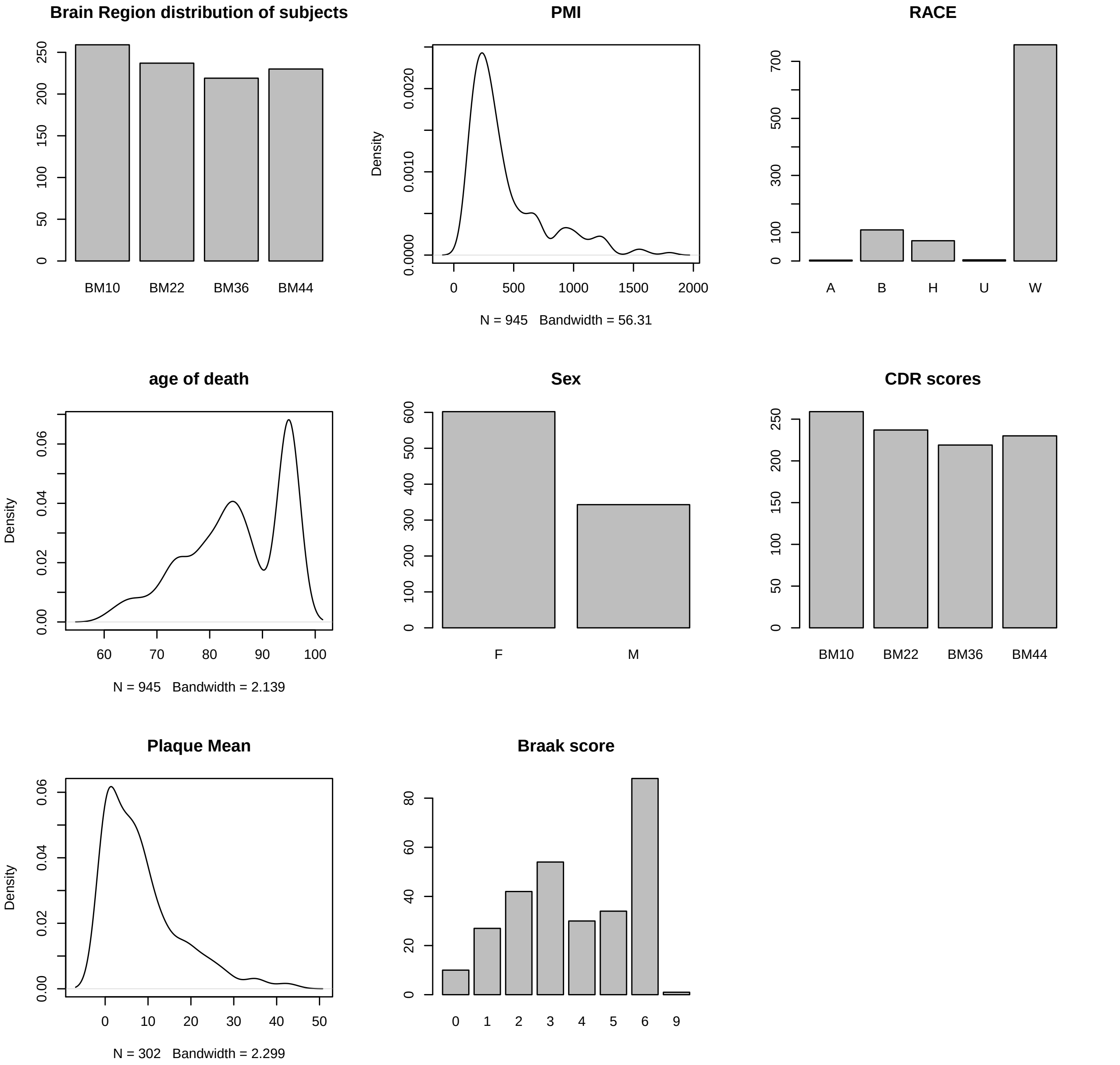


Figure S1: Clinical information of AD subjects. RNA-seq expression data for 945 autopsied samples from 364 subjects in four brain regions: frontal pole (BA10), superior temporal gyrus (BA22), parahippocampal gyrus (BA36) and frontal cortex (BA44). PMI: Post-Mortem Interval; NP.1: Neuropathology Category.

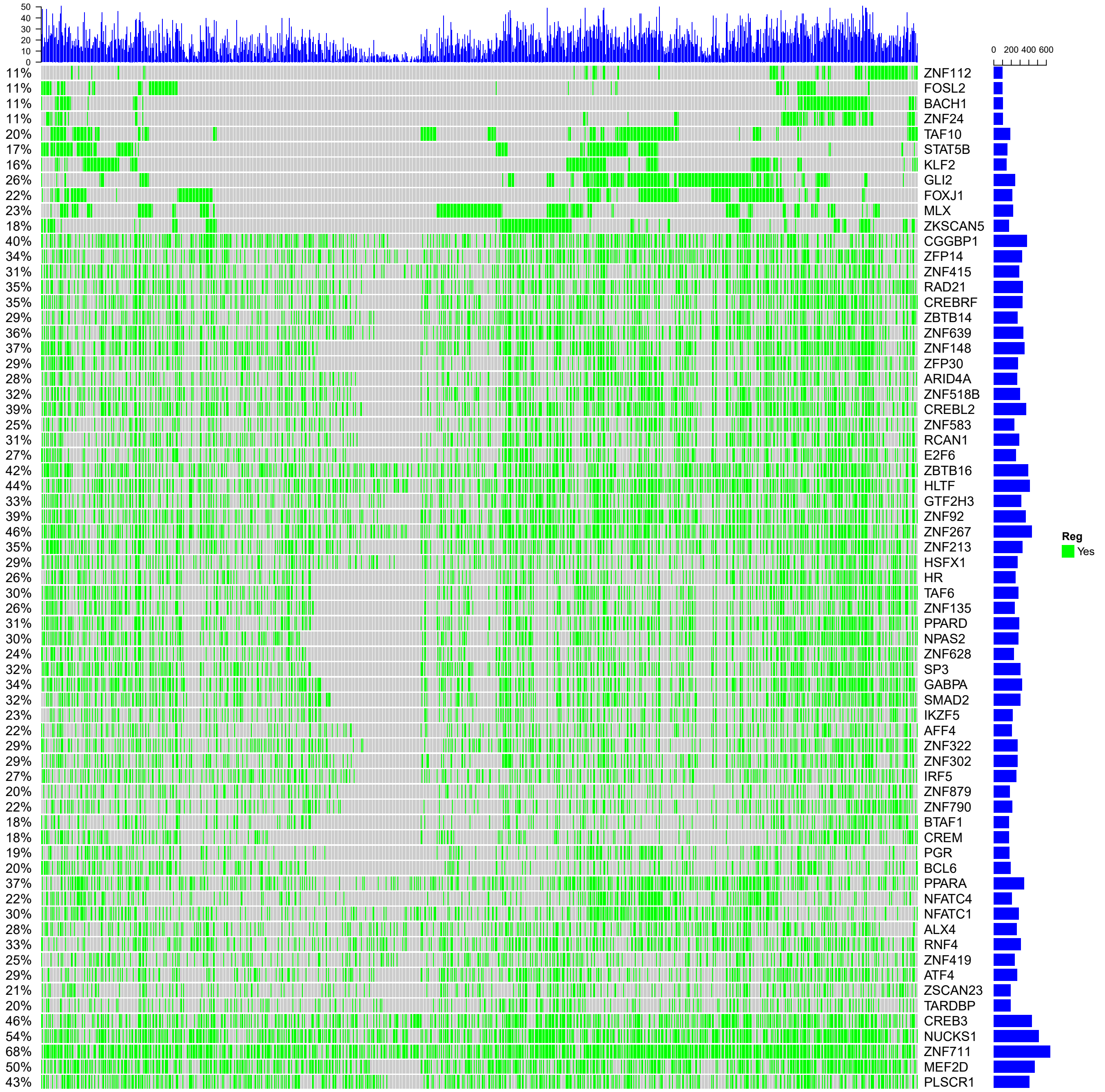


Figure S2: Regulation status of predicted regulators. Each bar in column indicates one subject and rows represents the regulation loss regulators. From this figure, we observe that TFs have different regulation status in selected subjects. This results suggests that transcription regulation loss is not specific to some AD patients but a general phenomenon during AD genesis and progression.

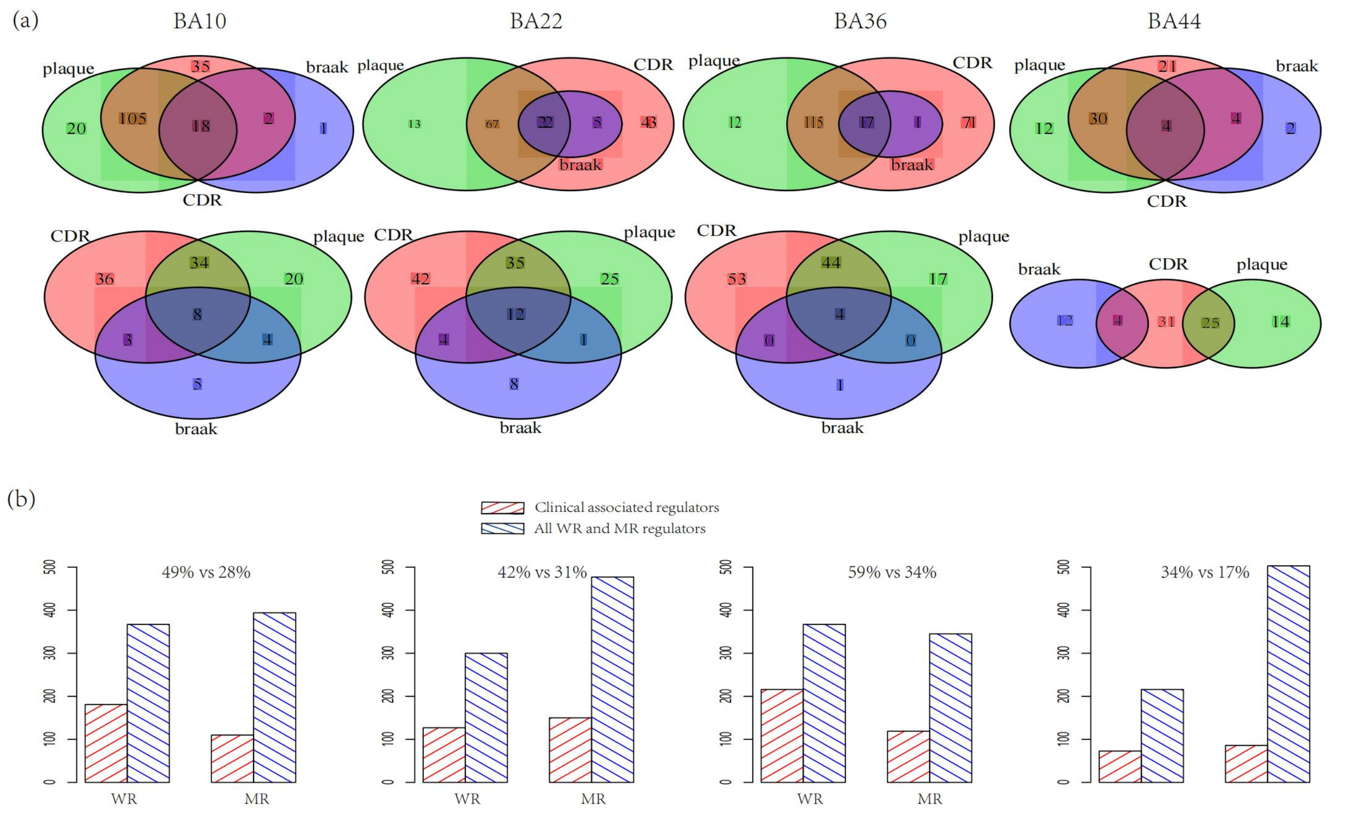


Figure S3: (a)The WR and MR regulators predicted with clinical association in four brain regions. Three clinical traits were used, including cognitive score (CDR), Braak score (braak) and amyloid plaque mean size (plaque). Four brain regions were studied: BA10, BA22, BA36 and BA44. (b)WR regulators more associated with clinical outcomes than MR. This is supported by Fisher's exact test, where the significance was $p = 6.01e-5, 3.8e-2, 8.32e-5 and 1.78e-4$ for BA10, BA22, BA36 and BA44, respectively.

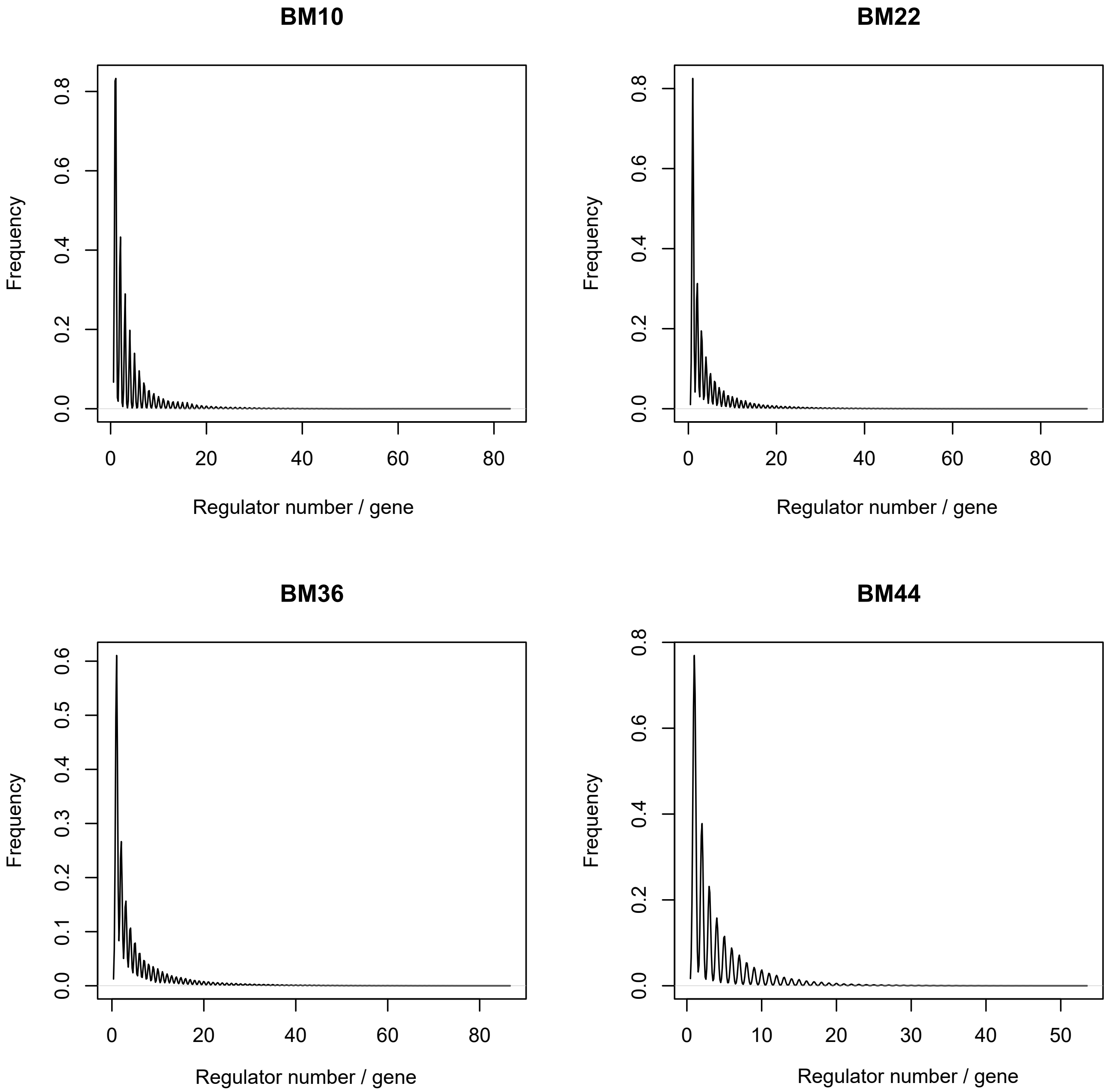


Figure S4: WR and MR regulators take dominant regulatory roles. 43% of genes are regulated by only one regulator and 80% are regulated by less than 5 regulators. Only very TFs take broad regulatory roles.

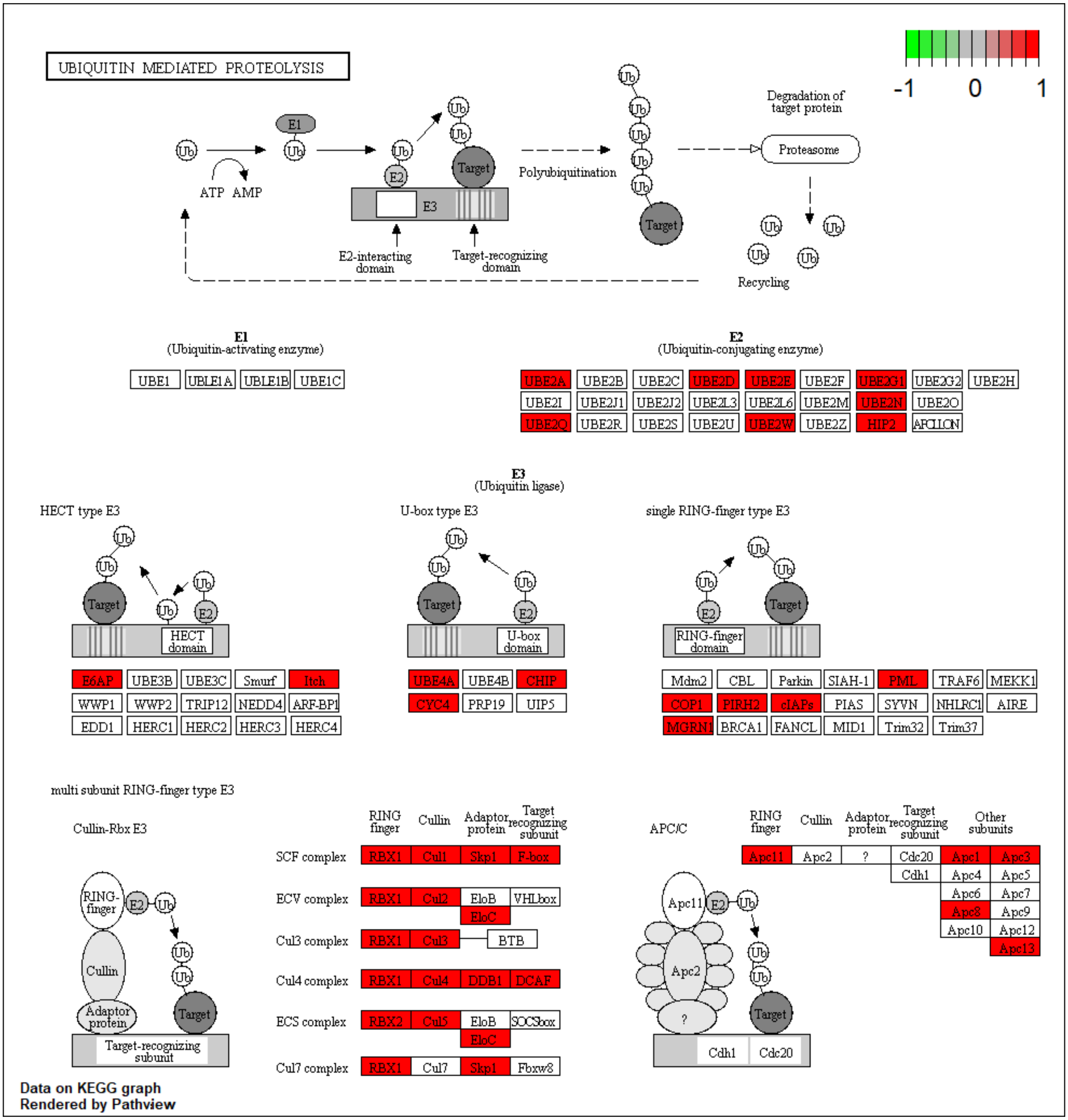


Figure S5: Ubiquitin-proteasome system is disturbed by transcriptional regulation loss. Here, the KEGG pathway “Ubiquitin Mediated Proteolysis” is marked for the dysregulated genes

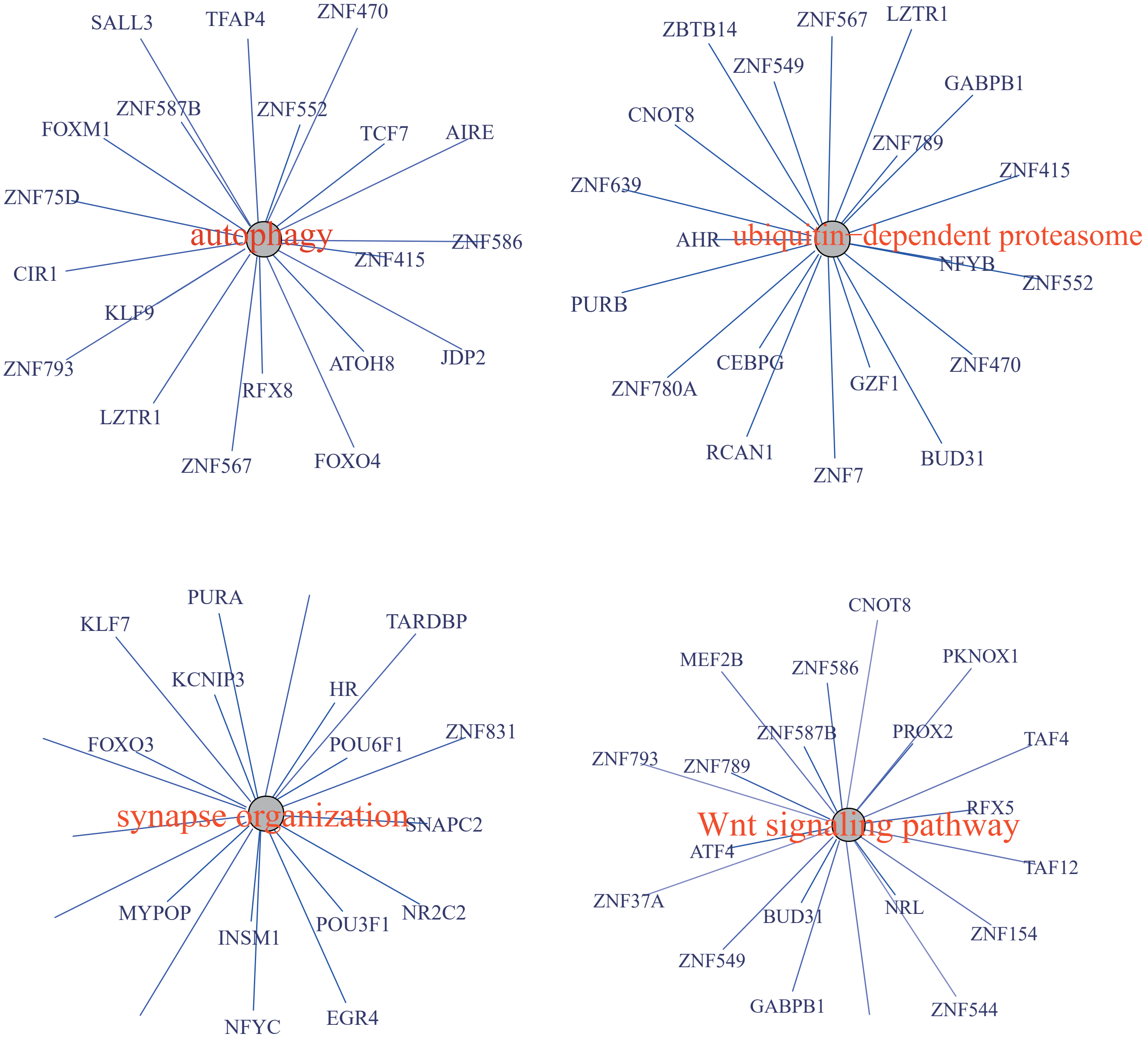


Figure S6: The top 20 predicted TFs associated with protein degradation processes, synapse function and Wnt signalling pathway.

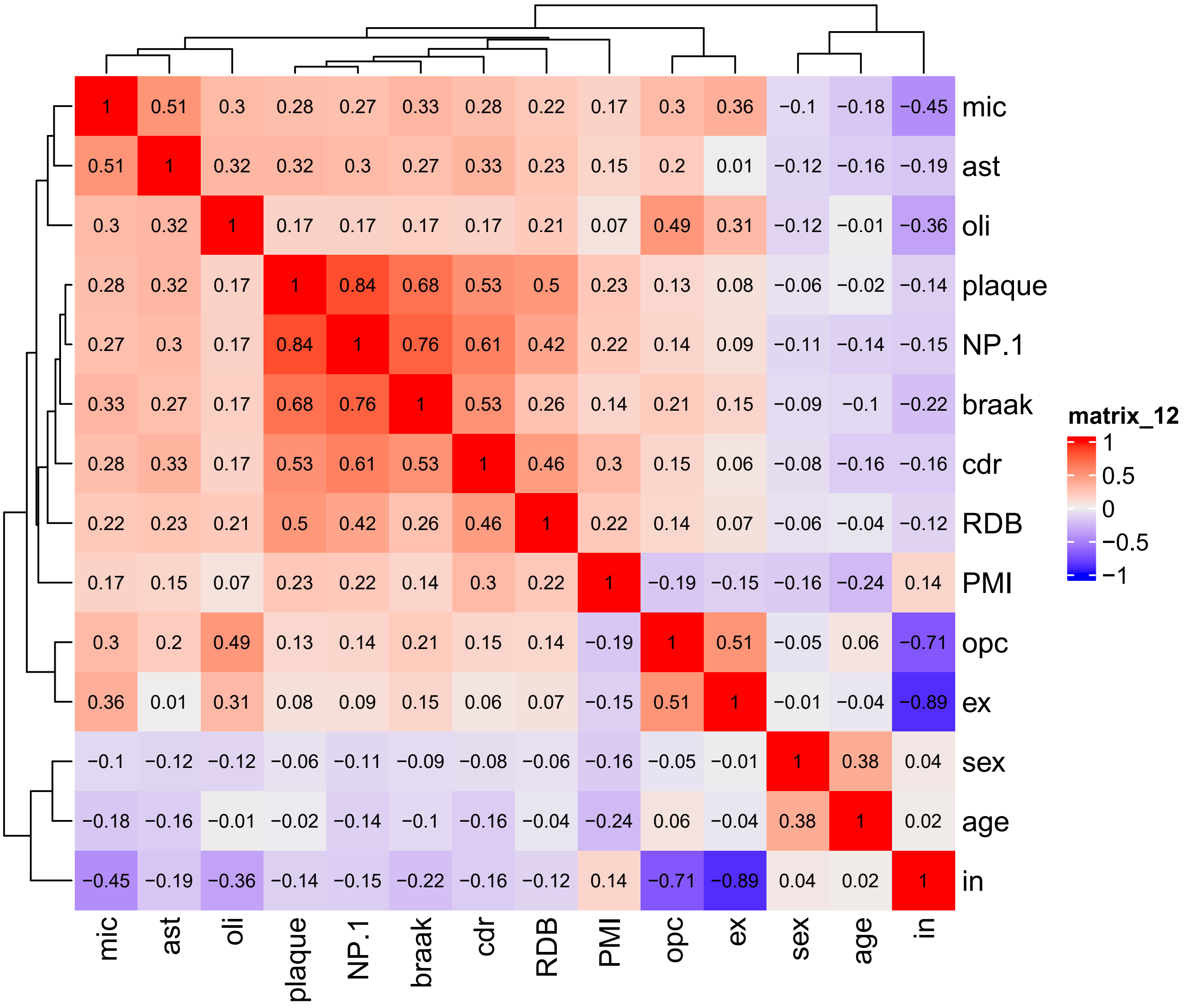


Figure S7: he correlation among different clinical traits and cell compositions

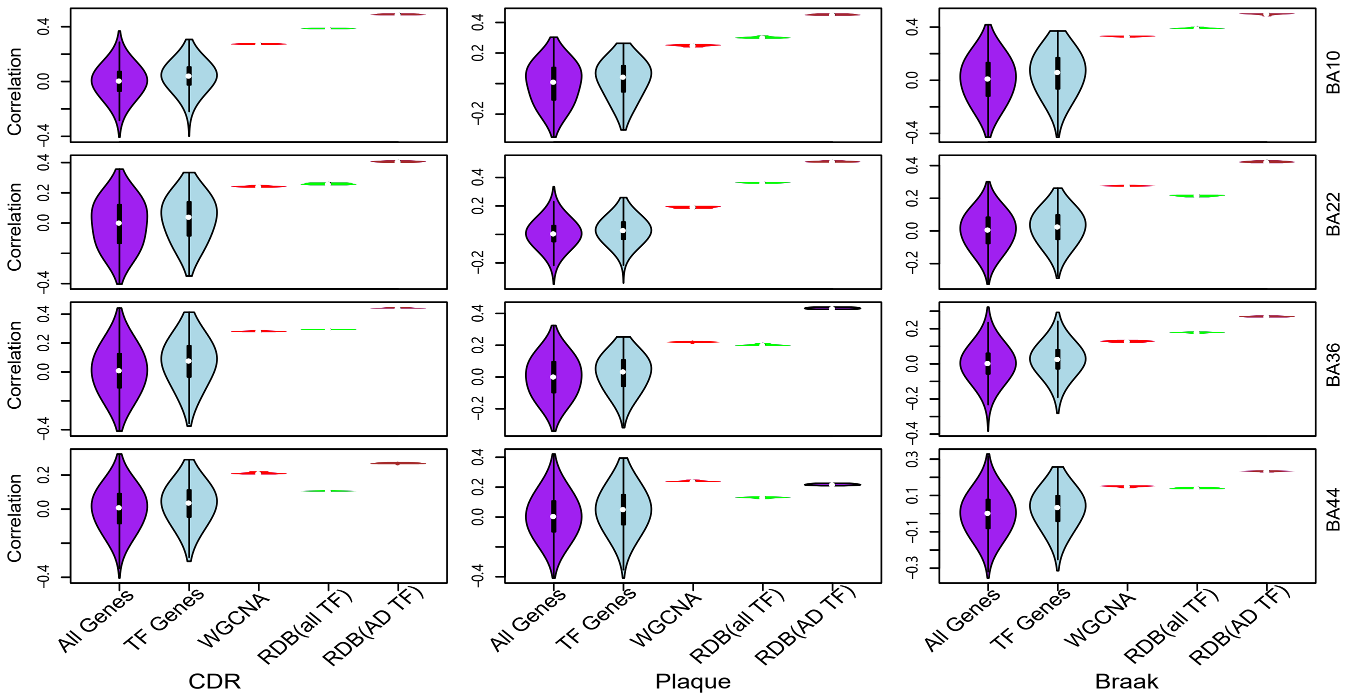


Figure S8: RDB better indicates clinical outcomes than genes, transcription factors and WGCNA modules. Here three clinical traits, including CDR, Plaque, Braak scores, were used.

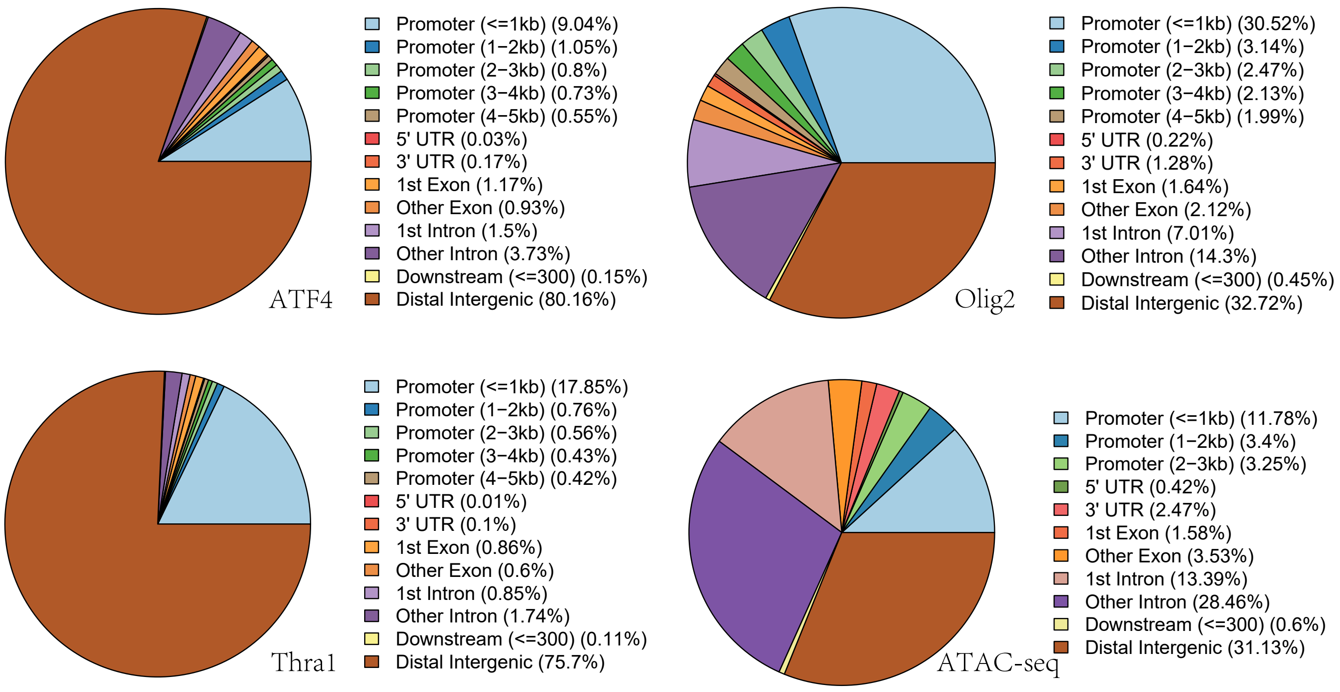


Figure S9: Genomic distribution of TF binding sites and ATAC-seq peaks.

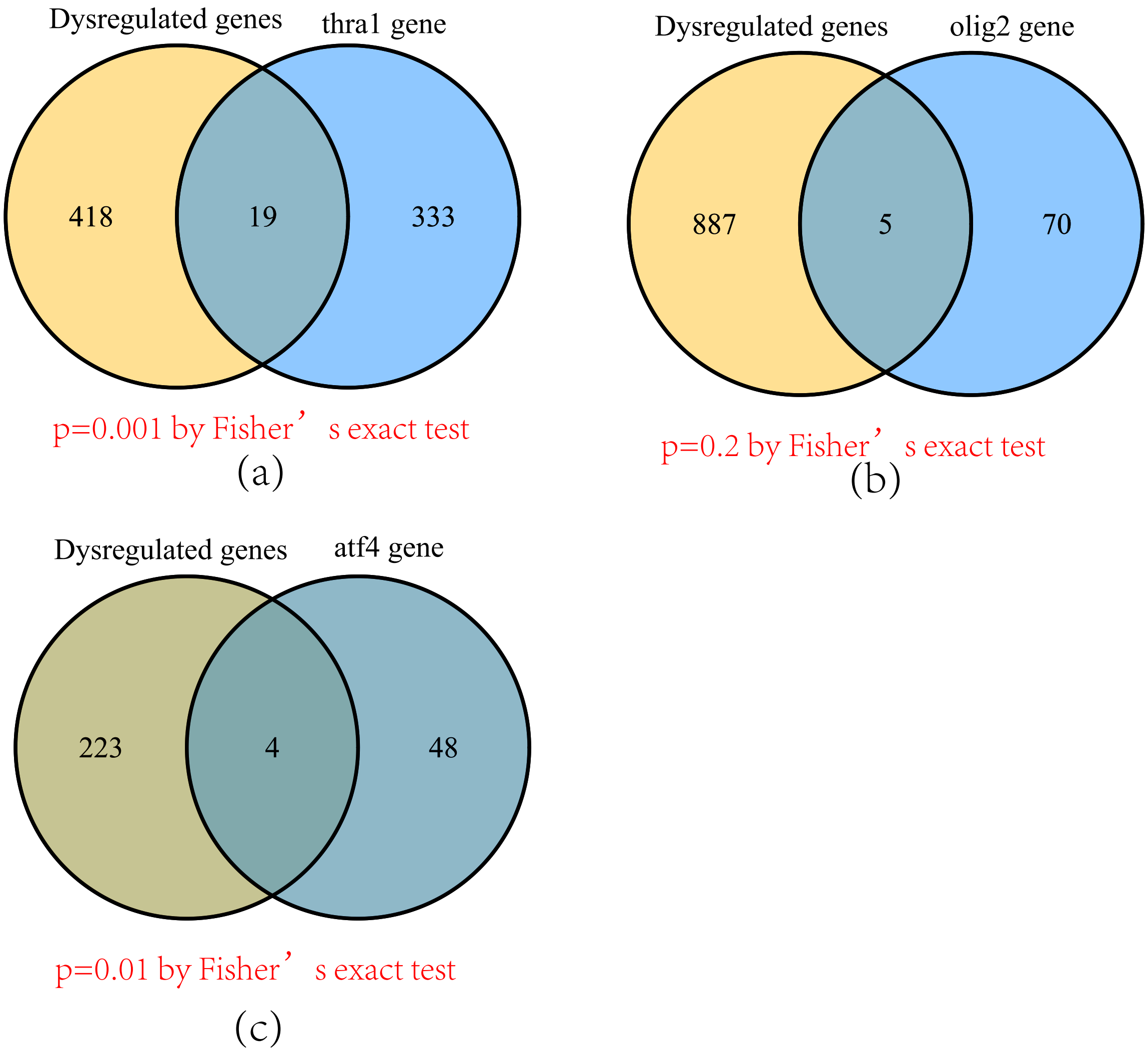


Figure S10: ChIP-seq peak loss of three TFs is overlapped with the predicted dysregulated genes. In this step, we collected the genes that were predicted to be regulated by (a) THRA1 (b) Olig2 and (c) ATF4. We checked their overlaps with the nearby genes of ChIP-seq peaks. Fisher's exact test was used to evaluate the significance. Significant overlaps were observed with THRA1 and ATF4. Here, only weak overlaps were observed, this might be caused by the many dysregulation TF regulation happened in the enhancer regions [3].

(a)

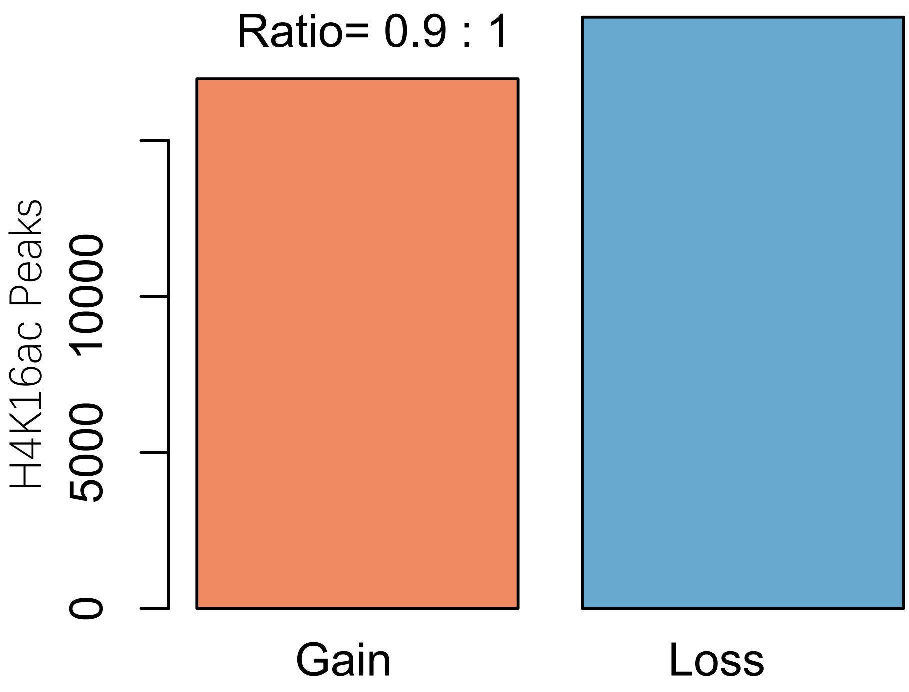


(b)

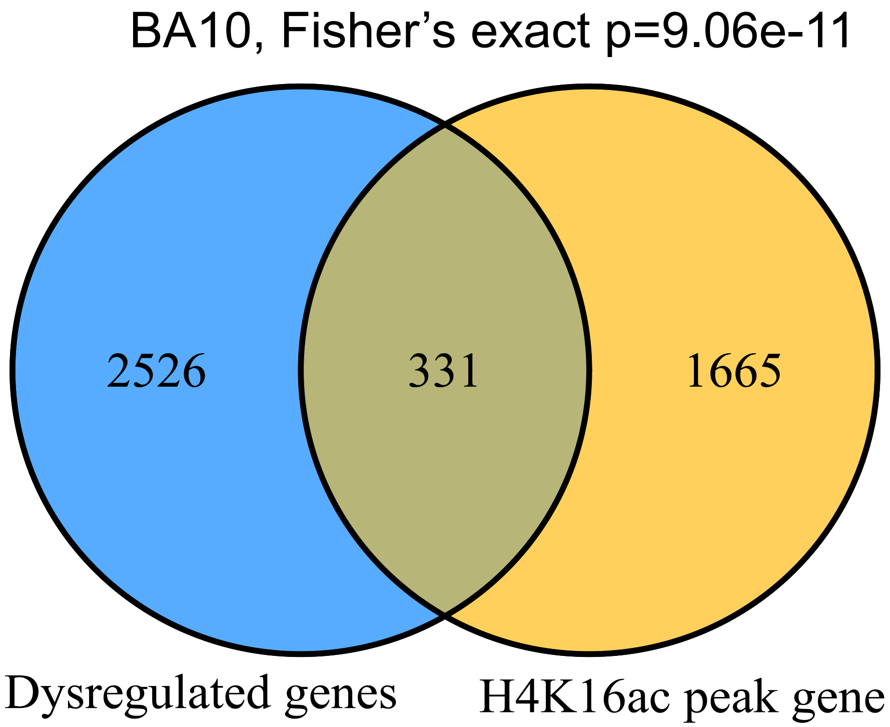


Figure S11: Regulation loss indicated by histone modification mark, H4K16ac. H4K16ac is a mark of active transcription and its ChIP-seq data were collected from published work [4]. (a) More regulation loss was observed based on H4K16ac peaks; (b) Regulation loss of H4K16ac peaks is greatly overlapped with the dysregulated genes identified in bi-clustering analysis.

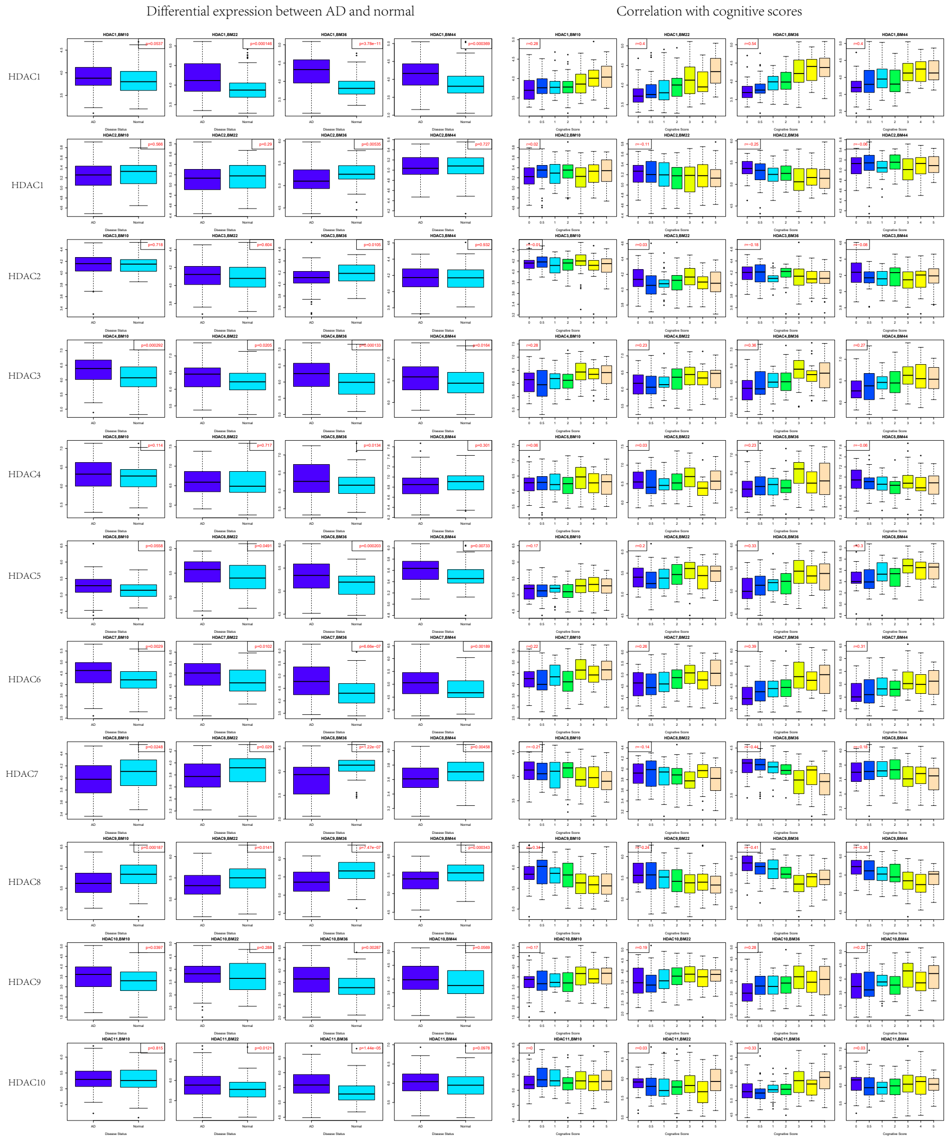


Figure S12: Differential expression and clinical association results of 11 HDAC genes

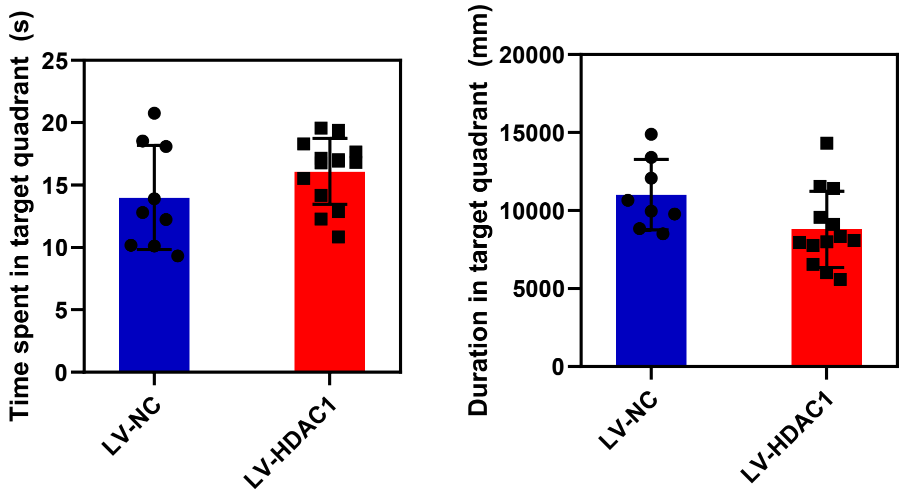


(a)

(b)

Figure S13: The water mirror maze results for (a) the spent time and (b) swimming distance in the target quadrant
